# Essential histone chaperones collaborate to regulate transcription and chromatin integrity

**DOI:** 10.1101/2020.11.04.368589

**Authors:** Olga Viktorovskaya, James Chuang, Dhawal Jain, Natalia I. Reim, Francheska López-Rivera, Magdalena Murawska, Dan Spatt, L. Stirling Churchman, Peter J. Park, Fred Winston

## Abstract

Histone chaperones are critical for controlling chromatin integrity during transcription, DNA replication, and DNA repair. We have discovered that the physical interaction between two essential histone chaperones, Spt6 and Spn1/Iws1, is required for transcriptional accuracy and nucleosome organization. To understand this requirement, we have isolated suppressors of an *spt6* mutation that disrupts the Spt6-Spn1 interaction. Several suppressors are in a third essential histone chaperone, FACT, while another suppressor is in the transcription elongation factor Spt5/DSIF. The FACT suppressors weaken FACT-nucleosome interactions and bypass the requirement for Spn1, possibly by restoring a necessary balance between Spt6 and FACT on chromatin. In contrast, the Spt5 suppressor modulates Spt6 function in a Spn1-dependent manner. Despite these distinct mechanisms, both suppressors alleviate the nucleosome organization defects caused by disruption of the Spt6-Spn1 interaction. Taken together, we have uncovered a network in which histone chaperones and other elongation factors coordinate transcriptional integrity and chromatin structure.

## INTRODUCTION

An enduring quest in the field of gene expression is to understand the function and coordination of the multitude of proteins required for transcription by RNA polymerase II (RNAPII). Distinct sets of proteins, often in complexes, dynamically associate with RNAPII during initiation, elongation, and termination (Kwak and Lis, 2013; Schier and Taatjes, 2020). Many of these proteins are required for transcription to occur on a chromatin template by helping to overcome the repressive effects of nucleosomes, by helping to maintain normal chromatin structure after the passage of RNAPII, or by the modification of histone proteins.

Histone chaperones, which comprise one class of factor essential for transcription, are a diverse set of proteins that directly modulate histone-DNA interactions in an ATP-independent fashion. In addition to transcription, histone chaperones are important during DNA replication and DNA repair (Hammond et al., 2017; Warren and Shechter, 2017). While the mechanisms and roles of some chaperones are well-understood (for example, (Chen et al., 2015; English et al., 2006; Liu et al., 2020)), the functions of most chaperones are not well defined (Hammond et al., 2017; Warren and Shechter, 2017). One mystery is why so many histone chaperones are required, as during transcription elongation, at least eight histone chaperones associate with elongating RNAPII. These include three that are conserved and essential for viability, have functional ties, and that are the focus of our studies: Spt6, Spn1/Iws1, and FACT.

Spt6 and Spn1 are two histone chaperones that directly interact with each other (Diebold et al., 2010; Li et al., 2018; McDonald et al., 2010; Reim et al., 2020). In addition, there is substantial evidence that their interaction is important for their functions in both yeast and mammalian cells (Diebold et al., 2010; McDonald et al., 2010; Yoh et al., 2008). However, there are several distinctions between Spt6 and Spn1. For example, in yeast, depletion of Spt6 results in massive changes in the specificity of transcription initiation on both the sense and antisense strands (Cheung et al., 2008; Doris et al., 2018; Gouot et al., 2018; Uwimana et al., 2017; van Bakel et al., 2013); in contrast, Spn1 has little effect on antisense transcription (Reim et al., 2020). Spt6 is required for the levels of certain histone modifications, including H3K36me2 and H3K36me3 (Carrozza et al., 2005; Chu et al., 2006; Gopalakrishnan et al., 2019; Youdell et al., 2008); in contrast, Spn1 is not needed for the level of these modifications, but it is required for their normal distribution on chromatin (Reim et al., 2020). Finally, while both Spt6 and Spn1 interact directly with histones (Bortvin and Winston, 1996; Li et al., 2018; McCullough et al., 2015), only Spt6 has been shown to interact directly with RNAPII (Sdano et al., 2017; Vos et al., 2018)., The association of Spn1 with RNAPII likely occurs indirectly, via Spt6 (Reim et al., 2020). Thus, both proteins clearly play vital roles, yet the nature of these roles and how they connect to each other is not well understood.

Spt6 also shares many functional similarities with FACT, another conserved and essential histone chaperone (Gurova et al., 2018). Both chaperones are believed to facilitate transcription of RNAPII through nucleosomes and/or to ensure nucleosome reassembly after the passage of RNAPII (for review see (Duina, 2011)). In addition, both regulate deposition of histone H2A.Z (Jeronimo et al., 2015). Furthermore, mutations in both cause similar effects on transcription (Cheung et al., 2008; van Bakel et al., 2013) and histone loss (Jeronimo et al., 2019). Despite their related functions, Spt6 and FACT clearly have independent, non-redundant roles, since both are essential for viability, as well as divergent biochemical activities, unique aspects of interactions with nucleosomes, and distinct patterns of recruitment to chromatin (Duina, 2011; Mayer et al., 2010; McCullough et al., 2015; Pathak et al., 2018). While one of the main mechanisms for recruitment of Spt6 to chromatin is by its interaction with RNAPII (Dronamraju et al., 2018; Mayer et al., 2010; Sdano et al., 2017), the recruitment mechanism for FACT is less clear and likely depends on multiple mechanisms, including interactions with histones (Cucinotta et al., 2019; Hodges et al., 2017), histone H2B ubiquitylation (Fleming et al., 2008; Murawska et al., 2020), and by the recognition of altered nucleosome structure (Martin et al., 2018). Thus, studies of Spt6, Spn1, and FACT have revealed intriguing relationships between the three and raised the question of how they interact during transcription.

In this work, we began by investigating the requirement for the Spt6-Spn1 interaction in *S. cerevisiae*, using *spt6-YW*, an *spt6* mutation that disrupts the Spt6-Spn1 interaction. We show that *spt6-YW* causes loss of Spn1 from the RNAPII transcription elongation complex and results in widespread defects in transcription and nucleosome positioning. By the isolation of extragenic suppressors of *spt6-YW*, we identify mutations in genes encoding several transcription elongation and chromatin factors. These include novel changes in FACT, as well as in Spt5, a conserved and essential elongation factor, revealing unexpected functional antagonism between these factors and Spt6/Spn1. By genetic and biochemical approaches, we show that the mechanisms by which alterations of FACT and Spt5 suppress *spt6-YW* are distinct, as altered FACT bypasses the requirement for Spn1 while suppression by altered Spt5 is Spn1-dependent. Despite acting through different mechanisms, both suppressors correct the chromatin defects of *spt6-YW*. Taken together, our studies reveal a network of interactions between essential histone chaperones and other conserved factors that is required for proper chromatin structure and gene expression.

## RESULTS

### The *spt6-YW* mutation impairs association of Spn1 with Spt6

To investigate the roles of the Spt6-Spn1 interaction, we used *spt6-YW*, a mutation predicted to impair the Spt6-Spn1 interaction due to the alteration of two conserved Spt6 residues located on the interface of Spt6 with Spn1 (Y255A, W257A) (Diebold et al., 2010; McDonald et al., 2010). Previously, *spt6-YW* was shown to cause strong mutant phenotypes, including temperature sensitive growth at 37°C, an Spt^−^ phenotype, indicative of transcriptional defects, and sensitivity to DNA damaging agents (Diebold et al., 2010).

To test whether *spt6-YW* affects Spt6-Spn1 interactions as predicted, we used co-immunoprecipitation analysis, comparing wild-type and *spt6-YW* strains. We immunoprecipitated Spt6 from cell extracts using a C-terminal triple-FLAG epitope tag and monitored co-immunoprecipitation of both Spn1 and Rpb1, the largest subunit of RNAPII. From wild-type extracts, we observed strong co-immunoprecipitation of both Spn1 and Rpb1 with Spt6, as expected (Figure 1A). In contrast, from *spt6-YW* extracts, we observed a dramatic decrease in Spt6-Spn1 co-immunoprecipitation, consistent with severe impairment of the Spt6-Spn1 interaction. However, the interaction of Spt6 with Rpb1 was largely unaffected by *spt6-YW*. We also tested *spt6-1004*, which contains a mutation outside of the Spt6 binding site for Spn1 (Kaplan et al., 2003), and found that this mutation did not impair either Spt6-Spn1 or Spt6-Rpb1 interactions. As the levels of Spn1 and Spt6 in the *spt6-YW* mutant extracts were comparable to wild-type levels (Figure 1A), the *spt6-YW* mutant phenotypes were likely caused by specific perturbation of the Spt6-Spn1 interaction rather than by altered protein levels. Thus, *spt6-YW* provides the opportunity to study the consequences of losing this conserved interaction between two essential histone chaperones.

**Figure 1.**
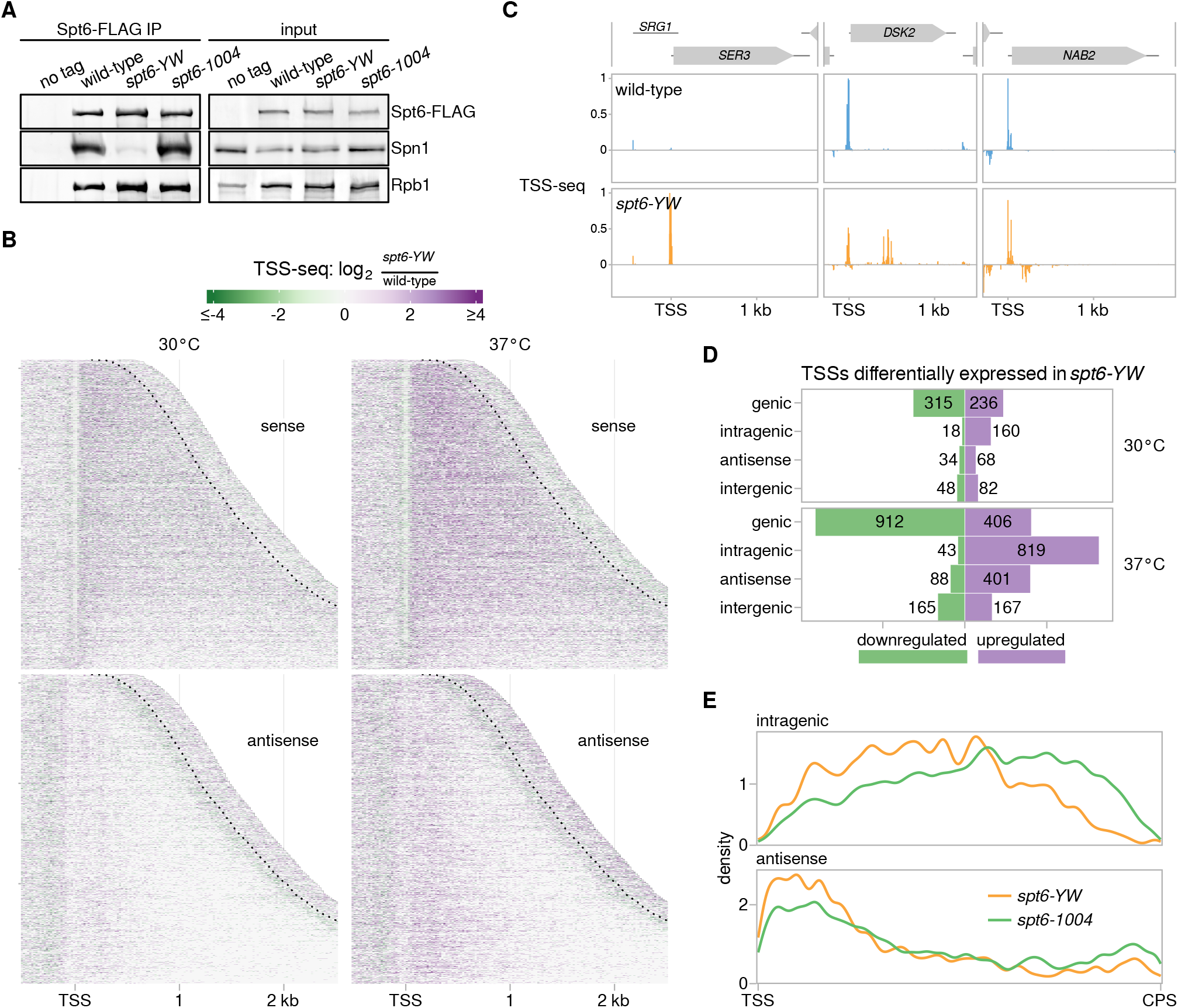
The *spt6-YW* mutation disrupts the Spt6-Spn1 interaction and causes altered sense and antisense transcription. (A) Western blots showing the levels of Spn1, Rpb1, and Spt6-FLAG in Spt6-FLAG immunoprecipitation (IP) samples and the corresponding inputs from untagged control (FY87), wild-type (FY3276), *spt6-YW* (FY3277), and *spt6-1004* (FY3283) strains. Spn1, Rpb1, and Spt6 were detected using anti-Spn1, 8WG16 anti-Rpb1, and anti-FLAG antibodies, respectively (see Methods). (B) Heatmaps of the ratio of TSS-seq signal in the *spt6-YW* (FY3223) strain over wild-type (FY87), grown at 30°C or with an 80 minute shift to 37°C. Data are shown for the sense and antisense strands of 3,087 non-overlapping verified coding genes aligned by wild-type genic TSS and sorted by length. The region shown for each gene extends up to 300 nt 3′ of the cleavage and polyadenylation site (CPS), which is indicated by the dotted line. (C) Examples of altered mRNA level (*SER3*), intragenic initiation (*DSK2*), and antisense initiation (*NAB2*) in *spt6-YW*. Relative TSS-seq signal in wild-type and *spt6-YW* strains shifted to 37°C is shown for each region, with sense and antisense signal plotted above and below the x-axis, respectively. Signal is independently scaled for each region shown. (D) Bar plots showing the number of TSS-seq peaks differentially expressed in *spt6-YW* versus wild-type, classified as described in Methods. “Intragenic” and “antisense" refer to sense-strand and antisense-strand intragenic TSSs, respectively. Green bars indicate downregulated peaks, and purple bars indicate upregulated peaks. (E) Distributions of the positions of upregulated intragenic TSSs on the sense (top) and antisense (bottom) strands along the scaled length of transcripts for *spt6-YW* (orange) and *spt6-1004* (green) at 37°C.

### The *spt6-YW* mutation alters transcription genome-wide

Previously we showed that Spt6 controls the fidelity of transcription initiation: when Spt6 was depleted using an *spt6-1004* mutant, transcription initiation decreased at thousands of genic transcription start sites (TSSs) and increased at thousands of intragenic TSSs on both the sense and antisense strands of genes (Doris et al., 2018). To study the contribution of the Spt6-Spn1 interaction to this function of Spt6, we performed transcription start site-sequencing (TSS-seq) (Arribere and Gilbert, 2013; Doris et al., 2018; Malabat et al., 2015) to quantitatively identify the 5′-ends of capped and polyadenylated transcripts across the genome at single-nucleotide resolution. This analysis was carried out for wild-type, *spt6-YW*, and *spt6-1004* strains grown at permissive temperature, 30°C, and after a shift to nonpermissive temperature, 37°C, for 80 minutes. We note that the shift to 37°C did not impair viability for either *spt6* mutant ((Doris et al., 2018); Methods).

Our TSS-seq results indicated extensive transcript level changes in *spt6-YW* compared to wild-type (Figure 1B, C), particularly with the shift to 37°C, after which over 1300 genic TSSs were misregulated and several hundred intragenic TSSs on both sense and antisense strands of genes were induced (Figure 1D). The transcriptional changes observed in *spt6-YW* at 37°C by TSS-seq were less severe than those observed in *spt6-1004* (Figure S1A-C), with the intragenic TSSs induced in *spt6-YW* in large part being a subset of those induced in *spt6-1004*: Of ~2600 sense strand intragenic TSSs induced in *spt6-1004*, about 29% were also induced in *spt6-YW*, alongside a small set of ~40 TSSs only induced in *spt6-YW* (Figure S1C). While sense strand intragenic TSSs induced in *spt6-1004* tended to occur towards the 3′ ends of genes as previously reported (Doris et al., 2018), the smaller set of sense strand intragenic TSSs induced in *spt6-YW* tended to occur towards the 5′ ends of genes (Figure 1E), suggesting that the Spt6-Spn1 interaction is involved in repressing a 5′-biased subset of intragenic transcripts. Antisense TSSs induced in both mutants showed similar preferences for the 5′ ends of genes (Figure 1E), similar to convergent antisense transcription observed in other cases (Kim et al., 2012; Lavender et al., 2016; Mayer et al., 2015; Shetty et al., 2017). Interestingly, the mean antisense TSS-seq signal in *spt6-YW* was greatest in the regions between the average wild-type positions of the +1, +2 and +3 nucleosome dyads (Figure S1D), suggesting a potential connection between these nucleosomes and antisense transcription.

As TSS-seq measures steady-state transcript levels, the induction of intragenic and antisense transcripts in *spt6-YW* could result from increased synthesis and/or reduced degradation of these transcripts. Previously, we showed that increased synthesis was the major contributor to induction of intragenic transcripts in *spt6-1004* (Doris et al., 2018). Since the intragenic and antisense transcripts induced in *spt6-YW* were largely a subset of those induced in *spt6-1004,* we expected the same to be true in the case of *spt6-YW*. To verify this, we performed native elongating transcript sequencing (NET-seq) to quantify elongating RNAPII in wild-type and *spt6-YW* strains. We then examined the NET-seq signal corresponding to antisense TSSs induced in *spt6-YW,* as antisense transcription is not obscured by overlapping genic transcription. We observed increased NET-seq signal in *spt6-YW* versus wild-type, consistent with increased synthesis contributing to the intragenic transcripts observed in *spt6-YW* (Figure S1E).

Previous studies showed that *spt6-1004* causes severely reduced levels of histone H3K36 di- and trimethylation (H3K36me2 and H3K36me3) (Carrozza et al., 2005; Chu et al., 2006; Gopalakrishnan et al., 2019; Youdell et al., 2008). As loss of the H3K36 methyltransferase Set2 causes increased levels of intragenic and antisense transcripts (Li et al., 2007; Venkatesh et al., 2016), this likely contributed to the transcriptional changes observed in *spt6-1004*. Therefore, we tested whether *spt6-YW* alters H3K36 methylation and found that, in contrast to *spt6-1004*, there was no significant effect on the total levels of H3K36me2 or H3K36me3 at either 30°C or after a shift to 37°C (Figure S1F). Thus, the widespread transcriptional changes and other phenotypes observed in *spt6-YW* were independent of changes in the level of these histone marks, although we have not excluded changes in the distribution of these histone modifications over genes.

### Suppressors of *spt6-YW* identify important transcription elongation factors

To gain greater insight into the requirement for the Spt6-Spn1 interaction, we conducted an unbiased selection for suppressors of the *spt6-YW* temperature sensitive phenotype (Figure 2A). We isolated 25 independent revertants of *spt6-YW* that allowed growth at 37°C and identified the causative mutations by a combination of whole-genome sequencing, complementation and linkage tests, and gene replacement. These analyses identified three classes of suppressors: extragenic (17 isolates, Figure 2A), intragenic (3 isolates), and strains disomic for chromosome 16 (8 isolates) (Table 1, Table S1).

**Figure 2.**
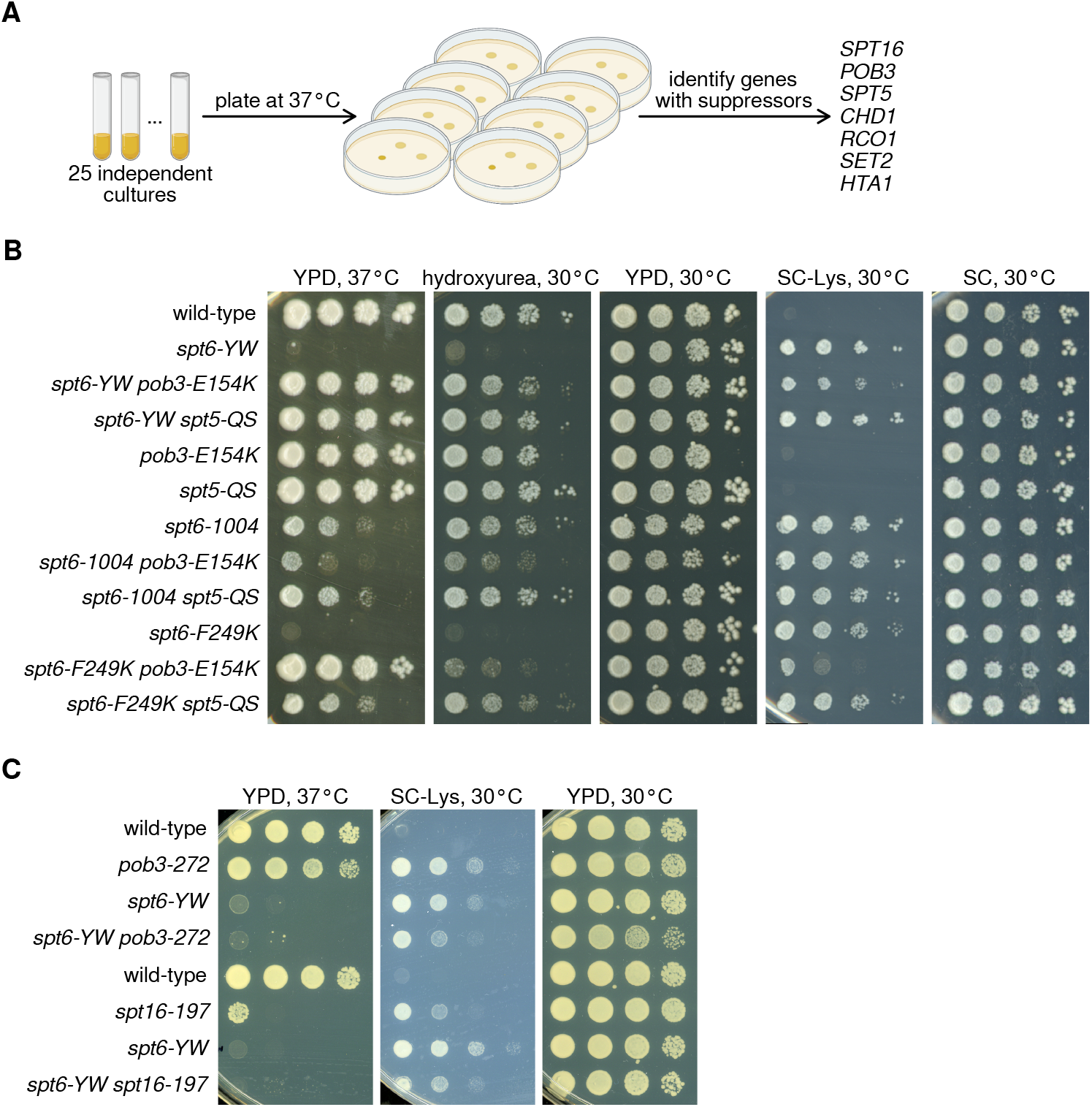
Mutations in *SPT16/POB3* and *SPT5* suppress *spt6-YW* in an allele-specific fashion. (A) A schematic showing the isolation of *spt6-YW* suppressors, with genes identified as extragenic suppressors listed on the right. (B, C) Analysis of genetic interactions between *SPT6*, *SPT5*, *POB3,* and *SPT16*. Strains were grown to saturation in YPD, serially diluted 10-fold, spotted on the indicated media, and grown at the indicated temperature.

While most of the analysis was done on the extragenic class, we first summarize our findings for the other two classes. The intragenic class consisted of three independent isolates that each contained a P231L amino acid change in Spt6 in addition to the original Y255A and W257A substitutions in Spt6-YW. P231 is located just outside of the region of Spt6 previously co-crystallized with Spn1 (Diebold et al., 2010; McDonald et al., 2010); the proximity of this position to the Spt6-Spn1 interface suggests that it suppresses *spt6-YW* by strengthening Spt6-Spn1 interactions. The eight disomic suppressors each contained an extra copy of chromosome 16, where *SPN1* is located. To test whether suppression was caused by the second copy of *SPN1*, we supplied *spt6-YW* mutants with an extra copy of *SPN1* on a centromeric plasmid and found that there was strong suppression of *spt6-YW* temperature sensitivity (Figure S2A). This result supports the idea that the major defect in the *spt6-YW* mutant is dependent on Spn1, consistent with previous studies (McDonald et al., 2010).

Analysis of the 17 extragenic suppressors identified mutations in seven genes, all of which encode factors that regulate transcription elongation and chromatin (Figure 2A). Four of these genes – *SET2*, *RCO1*, *CHD1*, and *HTA1* – were previously identified as suppressors of either *spt6* mutations or mutations that impair related elongation factors (Chu et al., 2006; Keogh et al., 2005; Lee et al., 2018; McCullough et al., 2015; Quan and Hartzog, 2010). As three of the genes, *SET2*, *RCO1*, and *CHD1*, are nonessential for viability, we tested complete deletions of each and found that they also suppressed *spt6-YW*, showing that suppression is conferred by loss of function (Table S2). The fourth gene, *HTA1,* is one of two genes encoding histone H2A, and the isolation of an *hta1* mutation is consistent with previous observations that amino acid changes in histones H2A and H2B suppress certain *spt6* mutations (McCullough et al., 2015).

In addition to these previously identified classes of suppressors, we identified novel mutations in three essential genes that have not previously been shown to suppress *spt6* mutations: *SPT16*, *POB3,* and *SPT5* (Table1, Table S1, Figure 2B). Spt16 and Pob3 are the two subunits of the histone chaperone FACT (Gurova et al., 2018), and Spt5 is a conserved elongation factor that interacts directly with RNAPII and has recently been implicated in controlling chromatin structure (Crickard et al., 2017; Ehara et al., 2019; Hartzog and Fu, 2013). The identification of *spt6-YW* suppressors in *SPT16*, *POB3*, and *SPT5* was unexpected, as previously isolated mutations in these genes conferred similar, rather than opposite phenotypes as *spt6* mutations (Cheung et al., 2008; Hainer et al., 2011; Hartzog et al., 1998; Jeronimo et al., 2019; Jeronimo et al., 2015; Kaplan et al., 2003; Lee et al., 2018; Malone et al., 1991; McCullough et al., 2015; Morillo-Huesca et al., 2010; Pathak et al., 2018; Swanson and Winston, 1992). The analysis of the FACT and Spt5 suppressors is the focus of the rest of these studies.

### Mutational changes in FACT and Spt5 suppress *spt6-YW* in an allele-specific fashion

Given the novel genetic suppression of *spt6-YW* by *pob3, spt16,* and *spt5* mutations, we asked whether the genetic interactions were allele specific. Allele specificity would suggest that suppression occurs by the alteration of specific molecular properties of these factors rather than by a general reduction of function. We tested this by constructing double mutants, combining *spt6-YW*, *pob3-E154K*, or *spt5-QS* with other mutations, and then assaying the double mutant phenotypes. In all cases, we observed strong allele specificity (Figure 2B,C and Table S2). First, we tested the ability of *pob3-E154K* and *spt5-QS* to suppress phenotypes of two other *spt6* mutations, *spt6-1004* and *spt6-F249K*. Unlike *spt6-YW*, the *spt6-1004* mutation does not impair Spt6-Spn1 interactions (Figure 1A); the temperature sensitive phenotype caused by *spt6-1004* became more severe with *pob3-E154K* and it was not significantly suppressed by *spt5-QS*. In contrast, *spt6-F249K*, which, like *spt6-YW*, does impair the Spt6-Spn1 interaction (McDonald et al., 2010), was suppressed by both *pob3-E154K* and *spt5-QS* (Figure 2B). Reciprocally, when *spt6-YW* was combined with previously isolated alleles of *pob3*, *spt16*, *spt4*, or *spt5*, we did not observe suppression of any *spt6-YW* phenotypes, and with *spt4Δ* and some *spt5* alleles we observed double mutant lethality (Figure 2C; Table S2). Taken together, these results demonstrate a highly specific set of genetic interactions between mutations that impair Spt6-Spn1 association and suppressors that alter FACT and Spt5.

### Biochemical and genetic evidence suggests that altered FACT bypasses the requirement for the Spt6-Spn1 interaction

Our genetic results suggest that the suppressor mutations might restore the Spt6-Spn1 caused by *spt6-YW*. To test this possibility, we conducted a series of co-immunoprecipitation experiments to assay interactions among five factors: Spt6, Spn1, FACT (Spt16), Spt5, and RNAPII (Rpb1 or Rpb3). These experiments were performed in wild-type and *spt6-YW* mutants with and without the suppressors. As an additional control, we included a *spn1-K192N* mutant which decreases the recruitment of Spn1 to chromatin (Zhang et al., 2008a). When we immunoprecipitated Spt6-FLAG from wild-type extracts, Spt16 and Spt5 both co-immunoprecipitated along with Spn1 and Rpb1, as expected (Krogan et al., 2002; Lindstrom and Hartzog, 2001; Lindstrom et al., 2003) (Figure 3A,B). In *spt6-YW* extracts, while there was a 20-fold decrease in Spn1 co-immunoprecipitation, the co-immunoprecipitation of Spt16, Spt5, and Rpb1 was not significantly altered (Figure 1A, 3A, and 3B). Reciprocal co-immunoprecipitation experiments strongly agreed with these findings and provided additional insight into the requirement of Spt6 for recruitment of Spn1 to RNAPII, based on the decreased signal in Spn1-Rpb1 and Rpb3-Spn1 co-immunoprecipitations observed in the *spt6-YW* background (Figure S2B and S2C). The total levels of the proteins assayed were not significantly changed in the *spt6-YW* mutant (Figure S2D-F). Together, these sets of co-immunoprecipitation results support the conclusion that *spt6-YW* specifically impairs the interaction of Spt6 with Spn1.

**Figure 3.**
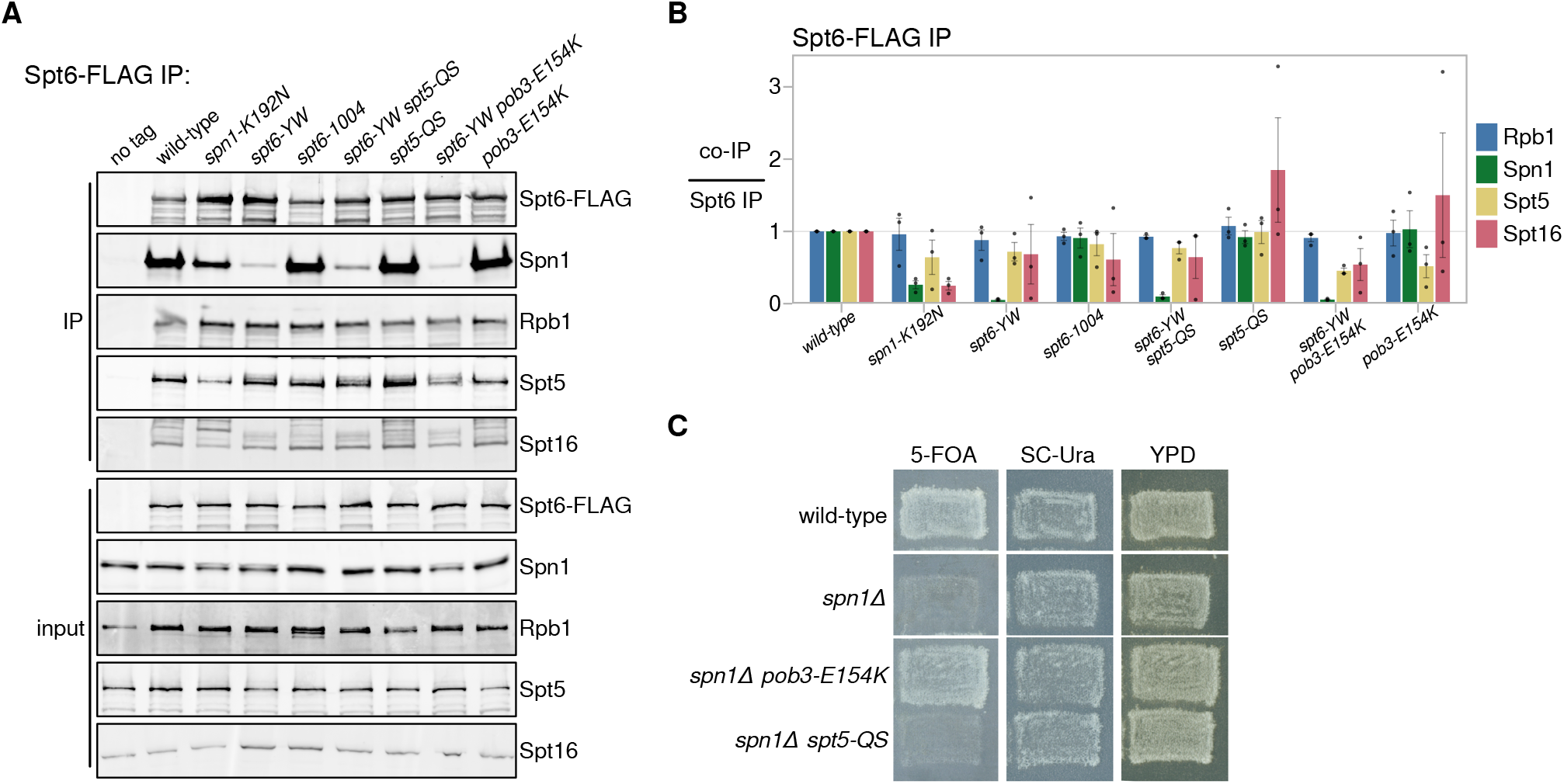
The *pob3-E154K* and *spt5-QS* suppressors do not restore the Spt6-Spn1 interaction. (A) Western blots for Spt6-FLAG co-immunoprecipitation analysis, as in Figure 1A. Spt5 and Spt16 were detected using their respective polyclonal antibodies. (B) Quantification of Spt6-FLAG co-immunoprecipitation experiments. Error bars indicate the mean ± standard error of the relative co-immunoprecipitation signal normalized to the Spt6-FLAG pull-down signal in the replicates shown. (C) Assay for the ability of *pob3-E154K* or *spt5-QS* to suppress *spn1Δ* inviability. Growth in the presence of 5-fluoroorotic acid (5-FOA) indicates viability after the loss of a *SPN1-URA3* plasmid as the sole source of Spn1.

Given the strong suppression of *spt6-YW* by *pob3-E154K* and *spt5-QS*, we were interested to test the effects of the suppressors on Spt6-Spn1 interaction. Our results showed that neither suppressor mutation altered the Spn1-Spt6 co-immunoprecipitation defect in an *spt6-YW* background (Figure 3A, 3B, and Figure S2B-F). We note that co-immunoprecipitation of either Spt5-QS or Spt16 from the *pob3-E154K* mutant with Spt6, Spn1, and RNAPII was not impaired, suggesting that the suppressor proteins interact at a normal level within the RNAPII elongation complex. From these results, we conclude that neither suppressor detectably restores the Spt6-Spn1 interaction.

Our co-immunoprecipitation results suggested that *pob3-E154K* and *spt5-QS* might bypass the requirement for the Spt6-Spn1 interaction. To test this idea, we assayed whether *pob3-E154K* or *spt5-QS* could suppress the inviability caused by a complete deletion of *SPN1 (spn1Δ*). Our results showed that the *spn1Δ spt5-QS* double mutant was inviable (Figure 3C and Table S2), indicating that suppression by *spt5-QS* is dependent on Spn1. In contrast, the *spn1Δ pob3-E154K* double mutant was viable and grew comparably to a wild-type strain (Figure 3C and Table S2), showing that the *pob3-E154K* mutation not only bypasses the Spt6-Spn1 interaction, but overcomes the requirement for Spn1, an essential histone chaperone. Combined with our co-immunoprecipitation results, we conclude that the *pob3-E154K* allele bypasses the requirement for Spn1 *in vivo*, while suppression by *spt5-QS* occurs by a different mechanism. We have focused the rest of our studies on understanding the mechanism by which *pob3-E154K* bypasses the need for Spn1.

### The RNAPII interactome is changed by the *spt6-YW* and the *pob3-E154K* mutations

To increase our understanding of the bypass of Spn1 by *pob3-E154K*, we expanded our physical analysis of how the *spt6-YW* and *pob3-E154K* mutations alter the RNAPII elongation complex. To do this, we immunopurified RNAPII complexes from wild-type, *spt6-YW*, *pob3-E154K*, and *spt6-YW pob3-E154K* strains, and identified the co-purified proteins by quantitative mass spectrometry (Methods). Similar to previous studies (Harlen and Churchman, 2017; Mosley et al., 2013; Tardiff et al., 2007), we identified 89 Rpb3-interacting proteins that were significantly enriched in at least one strain when comparing Rpb3 pull-down to a mock pull-down (Table S3).

The mass spectrometry data recapitulated our co-immunoprecipitation results for the association of Spn1 with Rpb3: Spn1 association was reduced 13-fold in *spt6-YW* compared to wild-type, and this decrease was not rescued in the *spt6-YW pob3-E154K* double mutant (Figure 4A,B). Though the greatest effect of *spt6-YW* on the RNAPII interactome was the depletion of Spn1, this was accompanied by other smaller-scale changes (Figure 4A,B), including reductions in the association of Elf1 and the TFIIF subunits Tfg1 and Tfg2, as well as increases in the association of the termination factors Rai1, Rat1, and Nrd1. These changes may be the consequence of loss of the Spt6-Spn1 interaction.

**Figure 4.**
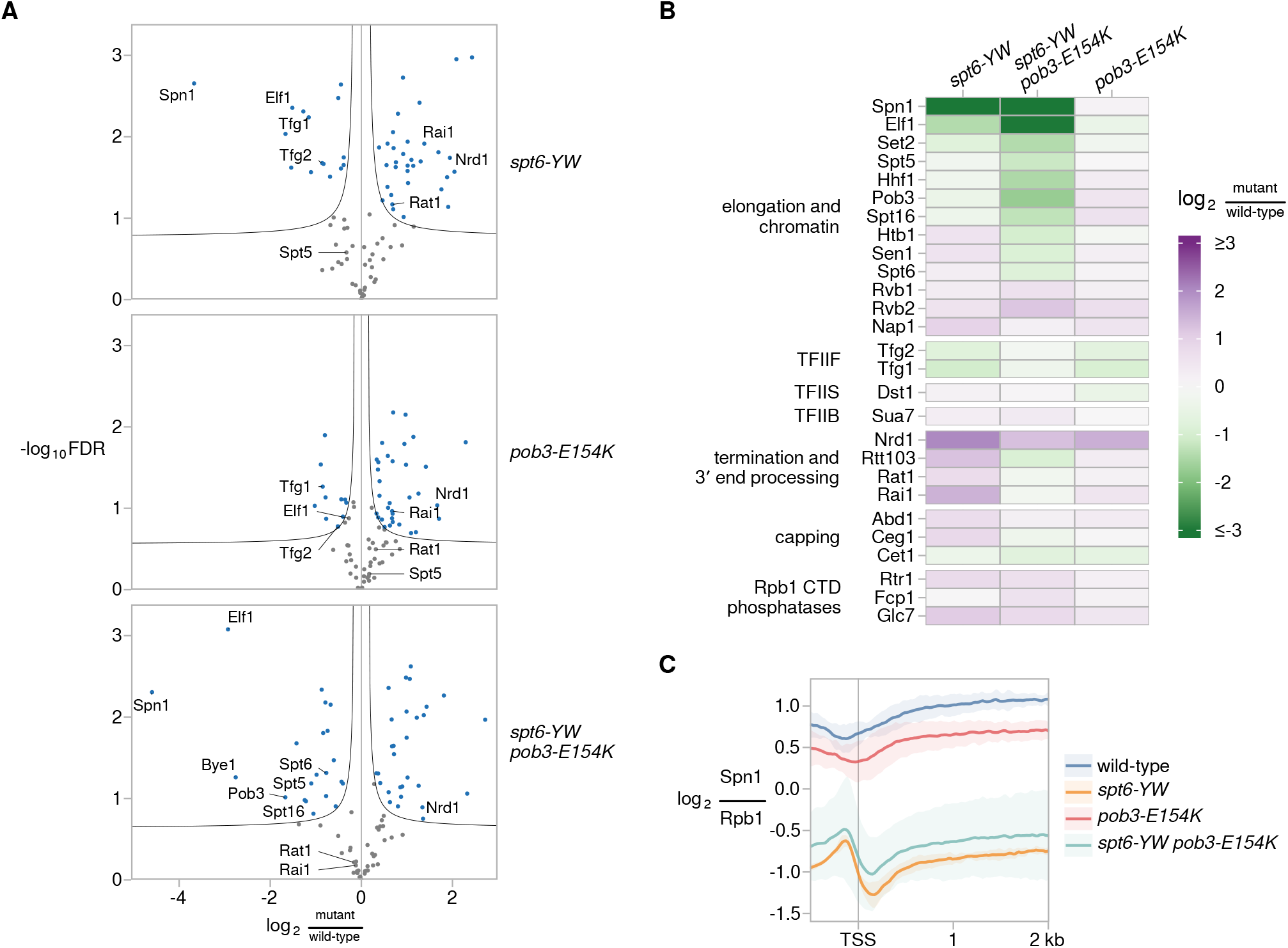
The *pob3-E154K* suppressor bypasses the Spt6-dependent recruitment of Spn1 to the elongation complex and chromatin. (A) Volcano plots comparing the Rpb3-FLAG interactome in *spt6-YW, pob3-E154K,* and *spt6-YW pob3-E154K* versus wild-type, as measured by mass spectrometry. Fold changes and significance values are calculated from Significance Analysis of Microarrays (Tusher et al., 2001), using either two (wild-type, *pob3-E154K, spt6-YW pob3-E154K*) or three replicates (*spt6-YW*). Black lines indicate significance cutoffs at an FDR of 0.1 and s_0_ of 0.1. Each point is an Rpb3-interacting protein enriched in Rpb3-FLAG IP samples over untagged mock IP samples, with blue points indicating proteins significantly changed between strains. (B) A heatmap of the ratio of mass spectrometry signal in *spt6-YW, spt6-YW pob3-E154K, and pob3-E154K* versus wild-type, for selected RNAPII-interacting factors. (C) The average Rpb1-normalized Spn1 ChIP enrichment over 3,087 non-overlapping verified coding genes aligned by TSS in wild-type (FY3292), *spt6-YW* (FY3289), *pob3-E154K* (FY3294), and *spt6-YW pob3-E154K* (FY3293). The solid line and shading are the mean and 95% confidence interval of the mean ratio over the genes considered from two replicates.

Interestingly, several changes to the RNAPII interactome were found specifically in the *spt6-YW pob3-E154K* double mutant and not in either single mutant. These changes included significantly reduced association of Spt6, Spt5, and both FACT subunits (Figure 4A,B). This suggests that decreased transcription elongation in *spt6-YW pob3-E154K* may be one mechanism by which this mutant overcomes loss of the Spt6-Spn1 interaction. Overall, our mass spectrometry results further support the model that *spt6-YW pob3-E154K* cells bypass the requirement for Spn1 and reveal changes in the RNAPII interactome that may allow this to occur.

### The Spt6-dependent recruitment of Spn1 to chromatin is not altered by *pob3-E154K*

Our co-immunoprecipitation results with *spt6-YW*, as well as recent results studying Spt6 depletion (Reim et al., 2020), suggest that Spt6 is required to recruit Spn1 to transcribed genes. Additionally, our co-immunoprecipitation results predict that such an impairment of Spn1 recruitment in *spt6-YW* would not be suppressed by the *pob3-E154K* mutation. To test these hypotheses, we performed chromatin immunoprecipitation sequencing (ChIP-seq) for Spn1 in wild-type and *spt6-YW* strains, with and without the *pob3-E154K* suppressor. To enable detection of global changes in Spn1 occupancy, we used exogenously added *S. pombe* chromatin for spike-in normalization, and to account for differences in Spn1 occupancy resulting from altered levels of transcription, we performed Rpb1 ChIP-seq from the same chromatin samples used for Spn1 ChIP-seq.

In wild-type cells, Spn1 was distributed over coding genes at levels highly correlated to levels of Rbp1, consistent with previous studies (Mayer et al., 2010; Reim et al., 2020) (Figure S3A). In *spt6-YW*, Spn1 occupancy over coding genes was decreased to about 18% of wild-type levels. This decrease in Spn1 occupancy in *spt6-YW* occurred uniformly over the length of RNAPII-transcribed genes, including virtually all protein-coding genes (Figure 4C and Figure S3B), snRNA genes, and snoRNA genes. In addition, the levels of Spn1 on chromatin in *spt6-YW* were not rescued by *pob3-E154K,* consistent with the bypass of Spn1 in that suppressor (Figure 4C and Figure S3B). Altogether, these results extend our co-immunoprecipitation results by showing that the Spt6-Spn1 interaction is required to recruit Spn1 to transcribed genes, and provide additional evidence that *pob3-E154K* bypasses the requirement for Spn1.

### The Pob3 and Spt16 suppressors are clustered in the FACT domain that interfaces with nucleosomal DNA

Our results showed that *pob3-E154K* alters FACT in a way that bypasses the requirement for *SPN1 in vivo*. To gain insight into how changes in FACT might cause this dramatic effect, we examined the locations of all the *pob3 and spt16* suppressors of *spt6-YW,* as well as eight additional *pob3* and *spt16* mutations that were isolated in a separate selection for suppressors of *spn1Δ* inviability, to be described elsewhere (F. López-Rivera, J. Chuang, R. Gopalakrishnan, D. Spatt, and F. Winston, unpublished). Most of the amino acid changes caused by these mutations are clustered within the dimerization domains of Pob3 and Spt16 (Figure 5A). We also note that many of the changes, including Pob3-E154K, are amino acid charge changes. The suppressor mutants are distinct in location from previously characterized *pob3* and *spt16* mutants that do not suppress *spt6-YW*, including *pob3-272* (I282K) and *spt16-197* (G132D) (Costa and Arndt, 2000; Feng et al., 2016; Jamai et al., 2009; Lycan et al., 1994; Malone et al., 1991; Rowley et al., 1991).

**Figure 5.**
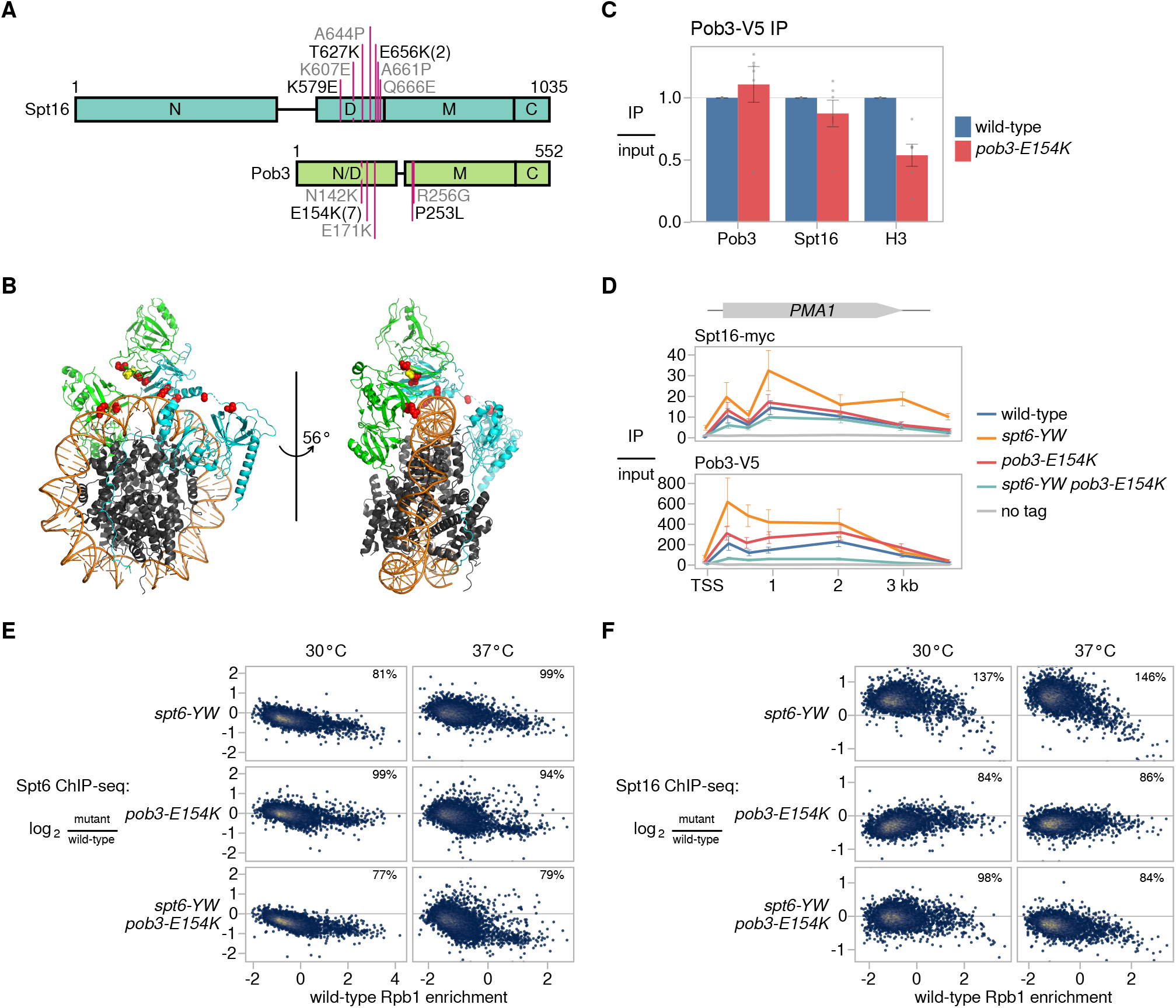
Evidence that mutant FACT suppresses the Spt6-Spn1 defect by restoring the balance between FACT and Spt6 on chromatin. (A) A schematic of the FACT subunits Spt16 and Pob3, depicting the amino acid changes caused by mutations suppressing *spt6-YW* (black) and *spn1*Δ (gray). The numbers in parentheses indicate the number of times a mutation was isolated if it was more than once. The labeled rectangles represent the domains of FACT: N – N-terminal, D – dimerization, M – middle, and C – C-terminal. (B) The structure of human FACT bound to a sub-nucleosomal particle (PDB: 6UPL, (Liu et al., 2020)), highlighting conserved residues corresponding to the locations of the yeast suppressor changes shown in (A). Human Spt16 (cyan), SSRP (green), and histones (grey) are shown as ribbon diagrams. The suppressor residues are shown as red spheres, except for SSRP1-E149, which corresponds to yeast Pob3-E154 and is shown as yellow spheres. Nucleosomal DNA is shown in orange. (C) Quantification of Pob3-V5 co-immunoprecipitation experiments. Error bars indicate the mean ± standard error of relative co-immunoprecipitation signal normalized to Pob3-V5 pull-down signal in the replicates shown. (D) ChIP analysis of Spt16 and Pob3 over the *PMA1* gene. The diagram shows Spt16 and Pob3 ChIP enrichment over input based on ChIP-qPCR measurements at the *PMA1* gene in wild-type, *spt6-YW*, *pob3-E154K*, and *spt6-YW pob3-E154K* strains. Error bars indicate the mean ± standard error of two replicates for each qPCR amplicon. (E) Scatterplots showing change in Spt6 ChIP enrichment in mutants (FY3277, FY3281, FY3282) over wild-type (FY3276) versus wild-type Rpb1 ChIP enrichment for 5,091 verified coding genes. Rpb1 enrichment values are the relative log_2_ enrichment of IP over input. (F) As in (E), but for Spt16 ChIP enrichment, using strains FY3299-3302.

Most of the residues changed in the suppressors are conserved between yeast and human, which allowed us to map their locations onto a published cryogenic electron microscopy structure of human FACT bound to a subnucleosome (Liu et al., 2020). Remarkably, the Spt16 and Pob3 residues targeted by the suppressor mutations map to the inner surface of the “saddle” module of FACT, which interfaces with nucleosomal DNA (Figure 5B). This structural analysis predicts that *pob3-E154K* would affect FACT-nucleosome interactions rather than Pob3-Spt16 interactions, despite their clustering in the dimerization domains. We tested this prediction by immunoprecipitating Pob3-V5 and assaying co-immunoprecipitation of Spt16 and histone H3. As predicted, co-immunoprecipitation of Spt16 with Pob3-V5 was unaffected by the *pob3-E154K* mutation, while co-immunoprecipitation of histone H3 with Pob3-V5 was decreased approximately two-fold (Figure 5C). These structural and biochemical results suggest that one mechanism by which *pob3-E154K* bypasses the requirement for Spn1 is by weakening the interaction between FACT and nucleosomes.

### Evidence that mutant FACT suppresses the Spt6-Spn1 defect by restoring the balance between FACT and Spt6 on chromatin

Elevated levels of either Spt6 or Spt16 cause mutant phenotypes, suggesting that the amount of Spt6 and FACT that is associated with chromatin is important for normal function (Clark-Adams and Winston, 1987; Malone et al., 1991). Based on this idea, as well as the suppression interactions between Spt6 and FACT, we tested whether the association of Spt6 and FACT with chromatin was altered in our mutants by performing ChIP-seq of Spt6 and Spt16 in wild-type and mutant strains. These experiments were done both at 30°C and after a shift to 37°C.

Our results showed that in most cases, the median recruitment of the mutant Spt6-YW protein was reduced to about 80% of wild-type Spt6 (Figure 5E and Figure S4A). The reduction in Spt6-YW occupancy was detected in both the *POB3* and the *pob3-E154K* backgrounds. In the one case where a reduction was not observed, the *spt6-YW* single mutant after a shift to 37°C, this was likely caused by the increased occupancy of RNAPII over gene bodies, particularly near 5’ ends (Figure S4B). Overall, these data suggest that *spt6-YW* caused a moderate decrease in Spt6 occupancy. This decrease is likely the result of the lack of Spn1 recruitment in this mutant (Reim et al., 2020).

In contrast to the decreased level of recruitment of Spt6-YW, our results showed that Spt16 recruitment was broadly elevated in the *spt6-YW* single mutant (Figure 5D and 5F). In the *spt6-YW pob3-E154K* double mutant, the recruitment of Spt16 was either restored to wild-type levels (at 30°C) or modestly decreased (at 37°C), suggesting that an increased level of FACT recruitment caused by *spt6*-YW was suppressed by *pob3-E154K*. The *pob3-E154K* mutation alone caused a moderate decrease in Spt16 occupancy over most genes, particularly after a shift to 37°C (Figure 5F, Figure S4A), consistent with the structural prediction and co-immunoprecipitation results that *pob3-E154K* causes a decreased association of FACT with nucleosomes.

We note that when we normalized Spt16 occupancy by Rpb1 ChIP-seq levels, we observed that the relative Spt16 occupancies are equivalent between wild-type and the *spt6-YW* mutant, suggesting that the altered recruitment of Spt16 in *spt6-YW* is coupled to changes in transcription (Figure S4A). From our data, we cannot distinguish whether increased FACT recruitment alters transcription or vice versa. However, regardless of the causal direction of this relationship, our data strongly suggest that *spt6-YW* causes increased recruitment of Spt16 and that this increase is suppressed by *pob3-E154K*. From these results we propose a model in which the levels of Spt6 and FACT on chromatin must be properly balanced for optimal function, and that an alteration in the level of chromatin association of Spt6 alters the level of association of FACT.

### Spt6, FACT, Spt5, and Spn1 functionally interact to modulate nucleosome organization *in vivo*

Spt6 is known to be required for normal nucleosome positioning and occupancy (DeGennaro et al., 2013; Doris et al., 2018; Ivanovska et al., 2011; Jeronimo et al., 2019; Jeronimo et al., 2015; Perales et al., 2013; van Bakel et al., 2013). To investigate the contribution of the Spt6-Spn1 interaction to this role of Spt6, we performed micrococcal nuclease sequencing (MNase-seq) on a set of wild-type and mutant strains grown both at 30°C and after a shift to 37°C. MNase-seq data from the wild-type strain showed the expected pattern over genes, with a nucleosome-depleted region upstream of the TSS and a regularly phased nucleosome array downstream (Figure 6A). In *spt6-YW*, the median distance between adjacent nucleosome dyads increased from the wild-type value of 165 bp to 169 bp at 30°C and to 171 bp at 37°C, manifesting as a progressive 3′ shift of the nucleosome array over genes (Figure 6A,B and Figure S5A). The *spt6-YW* mutation also caused increases in ‘nucleosome fuzziness’, i.e., the variability of nucleosome positions within the population, which we quantified as the standard deviation of dyad positions within each region occupied by a nucleosome. Median nucleosome fuzziness increased from 30.6 bp in wild-type to 32.5 bp in *spt6-YW* at 30°C, and from 30.6 to 34.6 bp at 37°C (Figure 6C). From these results, we conclude that the Spt6-Spn1 interaction directly or indirectly controls inter-nucleosome distance and variability in nucleosome positioning genome-wide.

**Figure 6.**
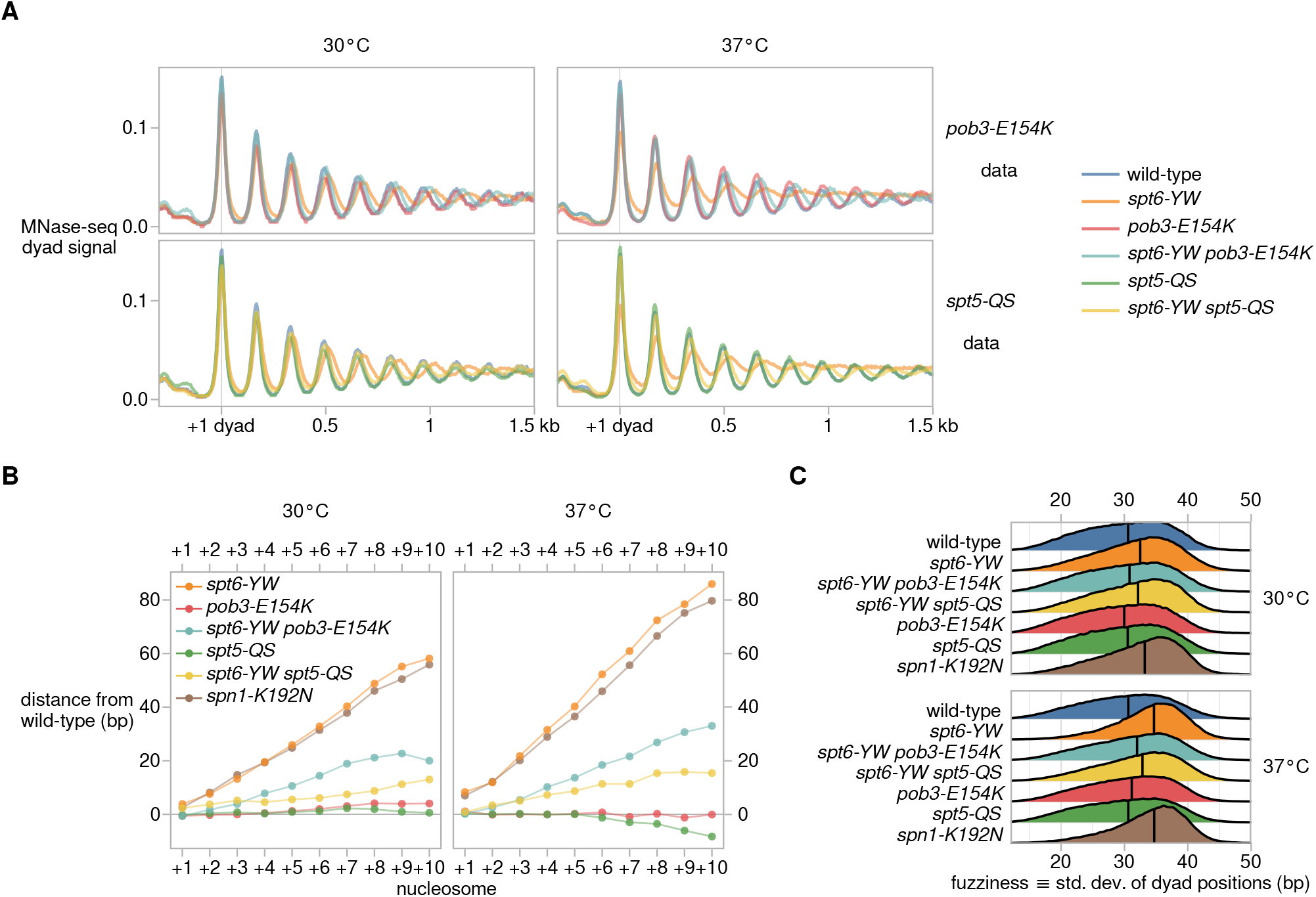
Spt6, FACT, Spt5, and Spn1 functionally interact to modulate nucleosome organization *in vivo*. (A) Average MNase-seq dyad signal over 3,086 non-overlapping verified coding genes aligned by 30°C wild-type +1 nucleosome dyad, for wild-type (FY87), *spt6-YW* (FY3223), *pob3-E154K* (FY3206) *, spt6-YW pob3-E154K* (FY3205) *, spt5-QS* (FY3273) *, and spt6-YW spt5-QS* (FY3274) strains grown at 30°C or with a shift to 37°C. Values are the mean of the mean library-size normalized coverage over the genes considered, over at least two replicates. (B) Mean differences in nucleosome position between mutant and wild-type, quantified from MNase-seq data. Nucleosomes are grouped based on their position in the nucleosome array of a gene, with the +1 nucleosome defined as the nucleosome region with midpoint position immediately 3′ of the TSS. (C) Distributions of nucleosome fuzziness, defined as the standard deviation of MNase-seq dyad positions within a nucleosome region. Vertical lines indicate median values of each distribution. Strains were grown at 30°C or with a shift to 37°C.

Strikingly, our MNase-seq results for *pob3-E154K* and *spt5-QS* indicated that each mutation suppressed the *spt6-YW* defects in inter-nucleosome distance and nucleosome fuzziness (Figure 6A-C and Figure S5A). The *spt5-QS* suppressor rescued *spt6-YW* inter-nucleosome distances to a greater degree than *pob3-E154K* (Figure 6B), while *pob3-E154K* rescued *spt6-YW* nucleosome fuzziness to a greater degree than *spt5-QS* (Figure 6C). Overall, the suppression of *spt6-YW* growth defects by *pob3-E154K* and *spt5-QS* correlates with suppression of the *spt6-YW* chromatin structure defects.

Finally, several results suggested that Spn1, like Spt6 and FACT, functions as a histone chaperone to regulate chromatin structure (Li et al., 2018; Reim et al., 2020); however, the role of Spn1 in global nucleosome organization has not yet been studied. Since *spt6-YW* greatly reduces the association of Spn1 with the RNAPII elongation complex, we hypothesized that loss of Spn1 might account for the nucleosome organization defects of *spt6-YW*. We investigated this possibility by studying the temperature-sensitive *spn1-K192N* mutant (Zhang et al., 2008a). MNase-seq of *spn1-K192N* revealed nucleosome positioning and fuzziness defects similar to those of *spt6-YW* (Figure 6B,C and Figure S5B). Additionally, the temperature sensitivity phenotype of *spn1-K192N* was suppressed by both *pob3-E154K* and *spt5-QS* (Table S2), hinting that the same mechanisms might suppress both *spt6-YW* and *spn1-K192N* chromatin defects. Thus, recruitment of Spn1 to the RNAPII elongation complex by Spt6 plays a significant role in the ability of both proteins to control chromatin structure across the genome.

## DISCUSSION

In this work, we have addressed the functional interactions of three essential and conserved histone chaperones, Spt6, Spn1, and FACT, during transcription in *S. cerevisiae*. We discovered that disruption of the Spt6-Spn1 physical interaction by *spt6-YW* impairs recruitment of Spn1 to the elongation complex. We went on to show that in the *spt6-YW* mutant, there are widespread changes in both transcription and chromatin structure, likely caused by the failure to recruit Spn1 to chromatin. Then, we identified suppressors of *spt6-YW,* revealing functional interactions with several regulators of chromatin and transcription. Focusing on two novel sets of suppressor mutations, in FACT (*SPT16* and *POB3*) and in *SPT5*, we showed that they suppress *spt6-YW* by distinct mechanisms. While our understanding of suppression by *spt5-QS* remains unclear, we demonstrated that a *pob3-E154K* suppressor mutation bypasses the need for Spn1. This bypass likely acts by creating an alternate chromatin state, possibly mediated by a weakened interaction between FACT and nucleosomes, which restores the balance between Spt6 and FACT on chromatin.

One striking result was the elevated level of FACT that was recruited to chromatin in an *spt6-YW* mutant observed in our ChIP-seq experiments, altering the normal ratio of chromatin-associated Spt6 and FACT. Interestingly, FACT levels and chromatin association are increased in cancer cells, consistent with the possibility that higher levels of FACT induce alterations in growth and transcription (Chang et al., 2018). We can imagine at least two possible scenarios by which the increase in FACT association might occur in *spt6-YW* mutants. The first model is based on the recent evidence that FACT associates with altered nucleosome structure (Martin et al., 2018). In this model, *spt6-YW* causes a disruption of normal chromatin structure, possibly due to loss of Spn1 recruitment, thereby increasing the level of the FACT-binding substrate, altered nucleosomes. In an alternative model, the increased levels of FACT are a consequence of defects in transcription caused by *spt6-YW* since FACT occupancy largely follows levels of Rpb1 (Feng et al., 2016; Martin et al., 2018). Although we cannot discern the cause and the effect relationship between chromatin and transcription from our current results, we favor the first possibility, due the novel nature of our suppressor mutations, which are predicted to weaken FACT-nucleosome interactions. Irrespective of the exact mechanism, our results are consistent with the concept that histone chaperones interact in a network that controls their relative levels on chromatin.

Suppression of *spt6-YW* by *pob3-E154K* mutation correlates with a reduction in the level of FACT association with chromatin (Figure 5F). Furthermore, during growth at 37°C, the levels of both chromatin-bound Spt6 and FACT are reduced in the *spt6-YW pob3-E154K* double mutant (Figures 5E, 5F, and S3B), yet the strain grows almost as well as wild-type (Figure 2B). This suggests that it is the Spt6:FACT ratio, rather than the absolute levels, that are critical for function. There is precedent for this among proteins that form structures, including bacteriophage heads (Floor, 1970; Sternberg, 1976) and histone proteins (Clark-Adams et al., 1988; Meeks-Wagner and Hartwell, 1986). As the functions of Spt6, FACT, and Spt5 are all sensitive to levels (Clark-Adams and Winston, 1987; Malone et al., 1991; Swanson et al., 1991), it seems plausible to speculate that the histone chaperones plus Spt5 function by a mechanism that requires a specific stoichiometry during RNAPII elongation.

We observed that in both *spt6-YW* and *spn1-K192N* mutants, inter-nucleosome distances are increased. Although we do not know the exact impact of this change on chromatin in our mutants, such a change might affect higher-order chromatin folding, chromatin compaction, and DNA accessibility (Correll et al., 2012; Li et al., 2016),possibly leading to some of the transcriptional defects observed in *spt6-YW*. Such an increase in inter-nucleosome distances has been observed in several other classes of mutants, including mutations that impair the chromatin remodelers Isw1, Isw2, Ino80, and Chd1 (Prajapati et al., 2020), as well as the Hir complex (Vasseur et al., 2016), a histone chaperone, and Yta7, a chromatin associated protein (Lombardi et al., 2011). Interestingly, we show that inactivation of one of these remodelers, Chd1, rescues *spt6-YW* and *spn1-K192N* mutant phenotypes (Table S2). At this point in our studies, we cannot distinguish whether Spt6 and Spn1 directly affect spacing as part of their interactions with histones, or whether this function occurs by coordination with other factors. Whatever the mechanism, it appears to be controlled by the same network of chaperones, as the change in the inter-nucleosome distances observed in *spt6-YW* is strongly suppressed by both the *spt5-QS* and *pob3-E154K* suppressors.

In conclusion, our studies have shed light on a network of interactions between histone chaperones that controls transcription and chromatin structure. Another recent study has shown that this network extends beyond Spt6, Spn1, and FACT to additional histone chaperones (Jeronimo et al., 2019). Given the essential and conserved nature of the three chaperones we studied, Spt6, Spn1, and FACT, it is surprising that complete loss of Spn1 can be strongly compensated by a single amino acid change in FACT. In spite of this, there must be strong selection to maintain this network of factors over evolutionary time, something that will be understood in greater depth as we learn the full range of the functions of these factors.

## Supporting information

Supplemental Figures and Tables for Viktorovskaya_2020

## ACKNOWLEDGMENTS

We thank Laurie Stargell for antisera, Tim Formosa for sharing yeast strains, plasmids, and antisera, Karolin Luger for helpful discussions, and Sarah Boswell and Mike Springer for use of their MiSeq. We also thank Karen Arndt and Catherine Weiner for helpful comments on the manuscript, and we thank Isabelle Washkurak for help with some of the experiments. Part of this research was conducted on the O2 High Performance Computing Cluster supported by the Research Computing Group at Harvard Medical School. This work was supported by NIH Fellowship F32GM119291 to O.V., a Ford Foundation Pre-Doctoral Fellowship to F.L.R., NIH Fellowship F31GM112370 to N.I.R., NIH Grant R01HG007173 to L.S.C., and NIH Grants R01GM032967 and R01GM120038 to F.W.

## AUTHOR CONTRIBUTIONS

O.V. and F.W. designed most of the experiments. O.V. performed the TSS-seq, ChIP-seq, MNase-seq, and co-immunoprecipitation experiments, and isolated and performed the genetic analysis of the *spt6-YW* suppressors. J.C. and D.J. performed and interpreted the bioinformatic analyses, with D.J. working under supervision of P.J.P. F.L.R. isolated and analyzed suppressors of *spn1Δ*. N.I.R. performed the western blots to measure protein levels in wild-type and mutant backgrounds and contributed to the genetic studies. M.M., F.W., and L.S.C. designed the NET-seq experiments and M.M. performed them. D.S. performed the single gene ChIP experiments and northern blots. O.V., J.C., and F.W. wrote the manuscript with feedback from all authors.

## DECLARATION OF INTERESTS

The authors declare no competing interests.

## STAR METHODS

### KEY RESOURCES TABLE

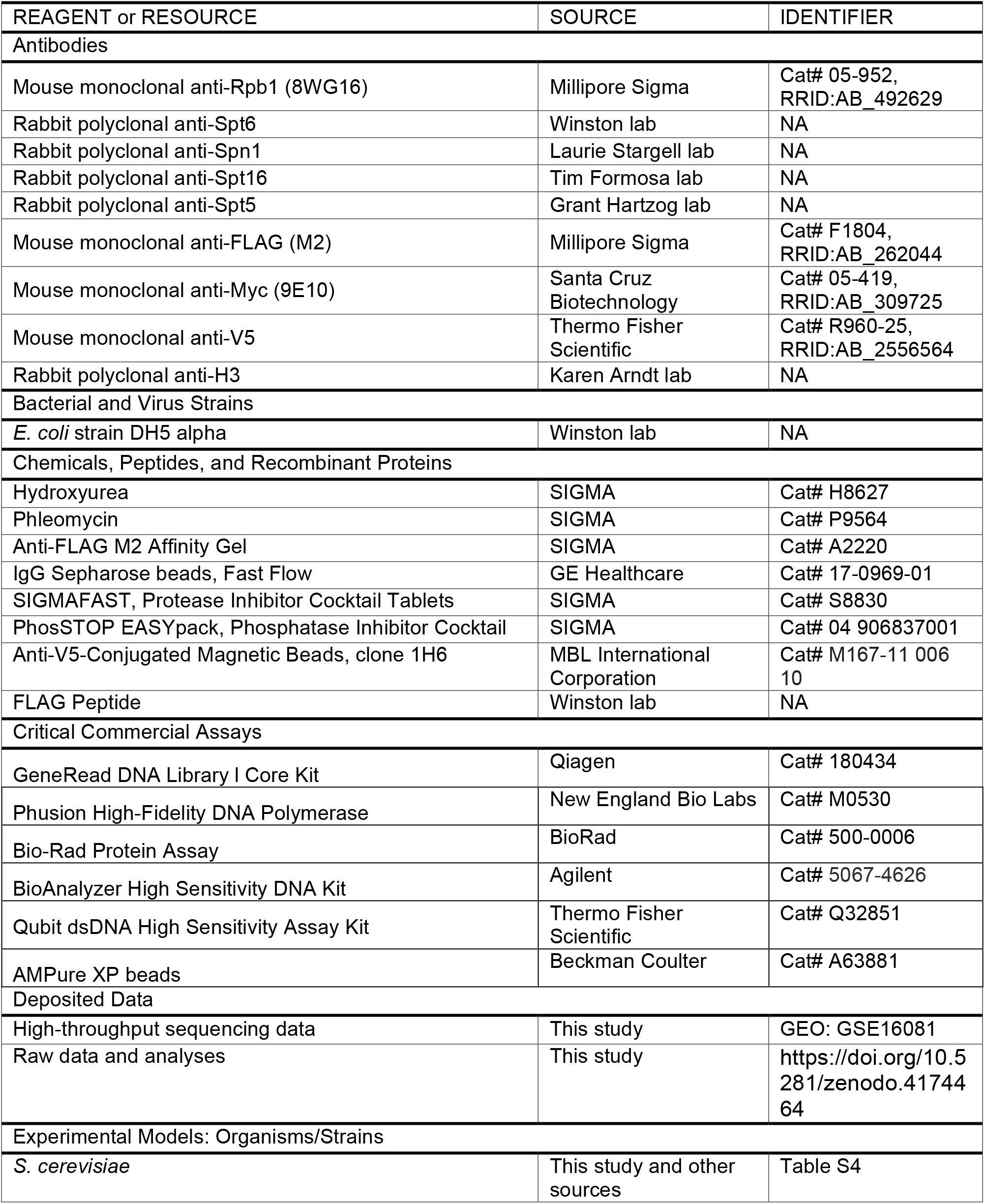
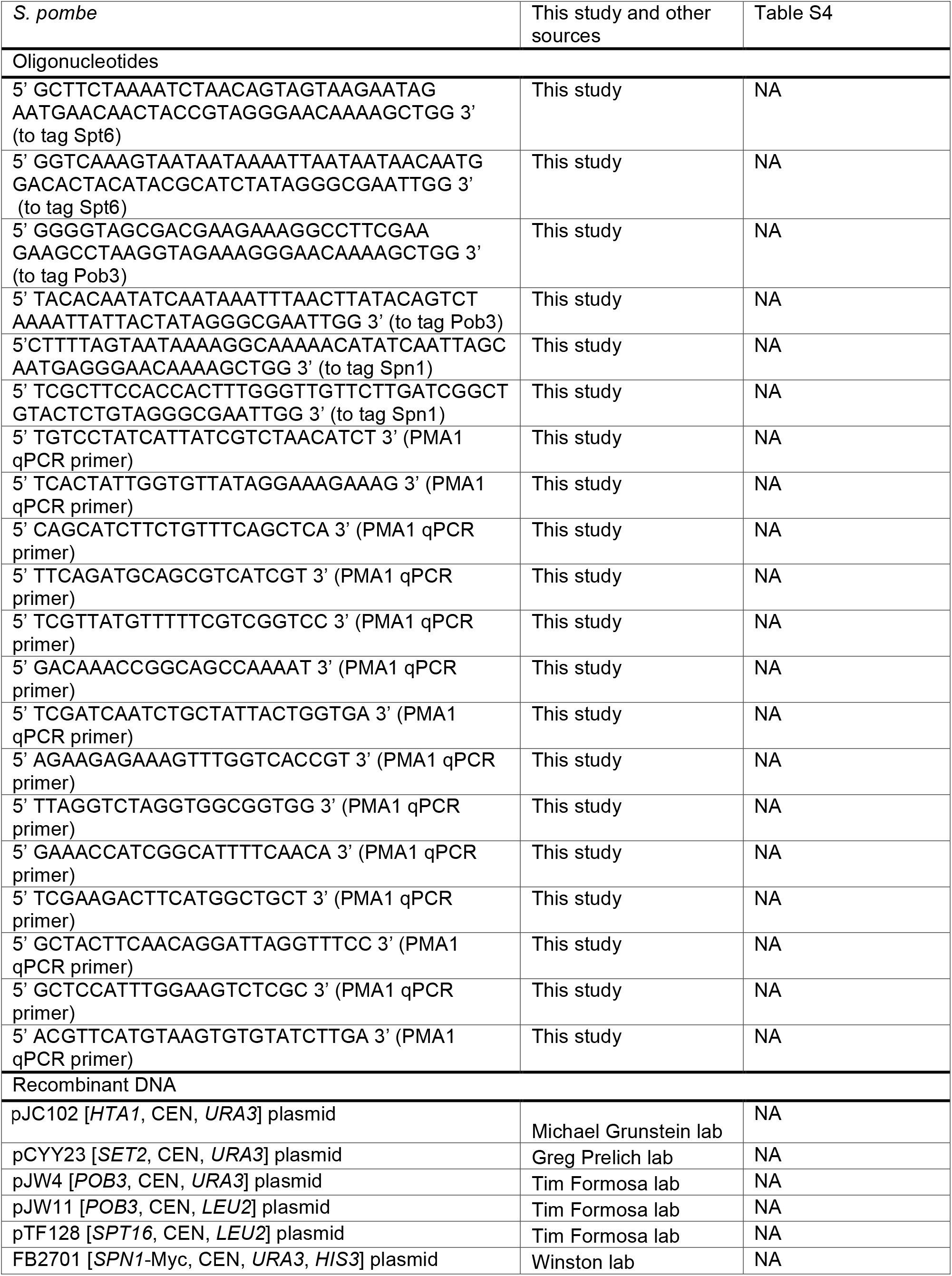
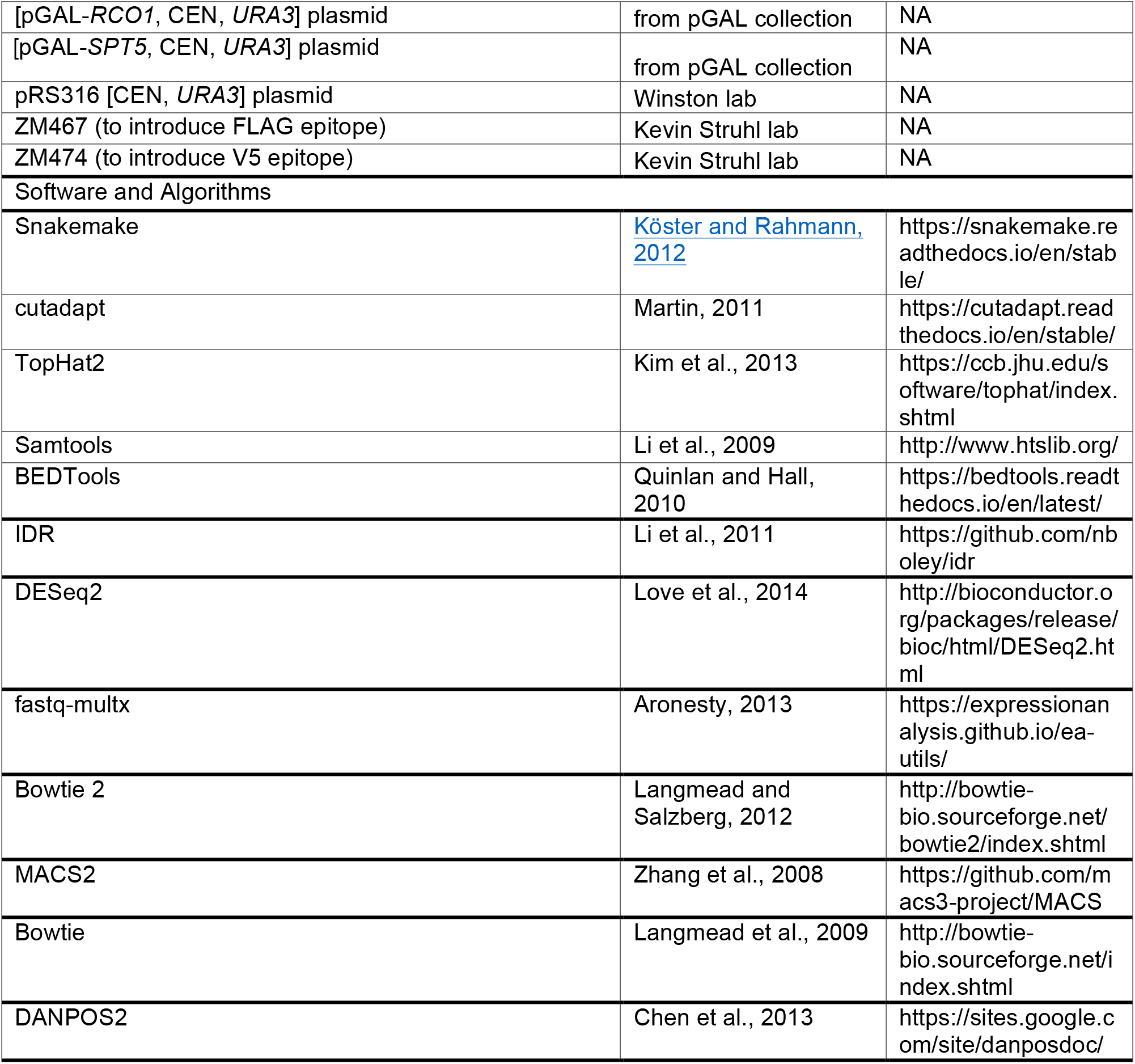

### RESOURCE AVAILABILITY

#### Lead Contact and Material Availability

Correspondence and requests for materials should be addressed to Fred Winston.

#### Data and Code Availability

All high throughput sequencing data except for whole genome sequencing data are available on GEO under accession number GSE160821. An archive containing code and raw data for reproducing all analyses except for whole genome sequencing and mass spectrometry analyses is available at Zenodo (https://doi.org/10.5281/zenodo.4174464). Additionally, updated versions of the Snakemake pipelines used are available at github.com/winston-lab.

### EXPERIMENTAL MODEL AND SUBJECT DETAILS

All experiments were performed in *Saccharomyces cerevisiae* S288C background (Winston et al., 1995). Strains used in this study are listed in Table S4. All strains were constructed by standard procedures, using either yeast transformation or crosses. The oligonucleotides and plasmids used for strain constructions are listed in the Key Resources Table. For spike-in normalization controls, we used *Schizosaccharomyces pombe.* For plasmid purification, we used *Escherichia coli DH5α*.

## METHOD DETAILS

### Yeast strains, media, and growth conditions

For TSS-seq, NET-seq, ChIP-seq and MNase-seq, yeast cultures were grown in YPD at 30°C or were shifted to growth at 37°C for 80 minutes before further processing or collection of the cells, as described previously (Cheung et al., 2008). The shift to 37°C for 80 minutes did not greatly affect the viability of the *spt6-YW* and *spn1-K192N* mutants resulting in 0.83±0.17 and 0.85±0.07 cell survival, respectively, compared to 30°C as measured by colony forming units (where CFU values for *spt6-YW* at 30°C relative to the wild-type is set as 1.00 and the SD from at least two independent measurements is indicated). To test yeast strains for sensitivity to hydroxyurea or phleomycin, YPD plates were supplemented with either hydroxyurea at a final concentration of 150 mM or phleomycin at 13 μg/ml, as previously described (Diebold et al., 2010). For genetic analysis, the double mutants were constructed using standard techniques.

### Co-immunoprecipitation

Yeast strains used for co-immunoprecipitation experiments are listed in Table S4. Cells were grown as 50-ml cell cultures to mid log phase (OD=0.6), collected, and lysed using buffer A [20 mM Hepes pH 7.6, 20% glycerol, 1 mM DTT, 1 mM EDTA, 125 mM potassium acetate, 1% NP-40 (IGEPAL, Sigma), 1 mM phenylmethylsulfonyl fluoride, 1X protease inhibitor cocktail (Sigma) and 1X phosphatase inhibitor cocktail (Sigma)] (Moqtaderi et al., 1996). For the FACT-histone co-IP experiments, lysates from strains FY3303-3306 were prepared using buffer B for lysis and coimmunoprecipitation [100 mM Hepes pH 7.9, 20% glycerol, 1 mM EDTA, 25 mM magnesium acetate, 0.4% NP-40 (IGEPAL, Sigma), 1 mM phenylmethylsulfonyl fluoride, 1X protease inhibitor cocktail (Sigma)]. Cell lysates were diluted with the lysis buffer to achieve equal protein concentrations (about 1 mg total protein per IP) and were then used for pull-downs with the respective antibody-conjugated beads: either the anti-FLAG M2-FLAG affinity gel (Sigma) (20 μl per IP) or anti-V5-conjugated magnetic beads (MBL International Corporation) (30 μl per IP) as previously described (Reim et al., 2020). The eluates were analyzed using western blotting. The antibodies used are listed in the Key Resource Table.

### Transcription start site sequencing (TSS-seq)

TSS-seq was performed as previously described (Doris et al., 2018). Analysis was done on yeast strains FY87, FY3223, and FY3125 after growth at 30°C or after a shift to to 37°C as described above. TSS-seq libraries were single-read sequenced on an Illumina HiSeq 2500 at the Harvard Bauer Core Facility.

### Native elongating transcript sequencing (NET-seq)

NET-seq was done as previously described (Churchman and Weissman, 2011). For strains FY2912 and FY3019, analysis was done on cultures grown at 30°C and after a shift to 37°C as previously described (Doris et al., 2018).

### Isolation and genetic analysis of *spt6-YW* suppressors

Yeast strains FY3019, FY3297, and FY3298, all containing the *spt6-YW* mutation, were used to isolate spontaneous and UV-induced suppressors of the temperature-sensitive (Ts^−^) phenotype as follows. Independent cultures were inoculated from single colonies and grown overnight to saturation in liquid YPD media. Then, 200 μl of each culture was plated on duplicate YPD plates, with one of them UV irradiated as previously described (Winston, 2008). The plates were incubated at 37°C and monitored daily for the appearance of colonies. Colonies were purified after either the third or fifth day of incubation at 37°C, yielding 52 independent suppressor candidates, and three colonies of each purified candidate was retested for suppression of the Ts^−^ phenotype. Genetic analysis was performed for the confirmed suppressor strains, excluding those which did not yield efficiently sporulating diploids. The remaining strains were screened to demonstrate a monogenic inheritance of suppression in a backcross to a parental *spt6-YW* strain to prioritize single-gene suppressor identification. This cross additionally allowed classification of suppressors as recessive, dominant, or intermediate based on the phenotype of the resulting diploids. Next, we performed linkage analysis, which revealed a total of three strains in which the suppressor phenotype was genetically linked to *SPT6*. Sanger sequencing of the open reading frame for *SPT6* from either of these independent suppressors uncovered same nucleotide change resulting in a P231L substitution, in proximity to the Y255A and W257A changes of the parental *spt6-YW* allele. Altogether, the genetic analysis resulted in identification of 25 strains, each presumably carrying an independent exogenous single-gene suppressor mutation summarized in Table S1.

### Identification of suppressor mutations

To identify causative mutations in the confirmed suppressor strains, we performed pooled linkage analysis, followed by whole genome sequencing (WGS) (Birkeland et al., 2010). Briefly, segregants with the suppressor phenotype (Ts^+^) were isolated from crosses between a suppressor strain (*spt6-YW sup*) and a parental *spt6-YW* strain, were pooled at equal cell number (12-50 segregants per pool), and sequenced. As a negative control, we included the parental strains as well as individual pools comprised of segregants with non-suppressor phenotype (Ts^−^) from backcrosses of three suppressors. DNA from each pool was extracted and used to generate genomic libraries as previously described (Gopalakrishnan and Winston, 2019). Single-read sequencing was performed either using an Illumina MiSeq according to the manufacturer’s instructions, or on an Illumina HiSeq 2500 by the Harvard Bauer Core facility. In parallel, while the WGS results for a subset of suppressors were emerging, we used linkage analysis and plasmid complementation to screen the remaining suppressors for mutations in already identified genes. This screen revealed that five suppressors likely contained mutations in *POB3*, which was further validated by Sanger sequencing identifying a *pob3-E154K* allele in each of these independent suppressors. The WGS data were processed using a custom pipeline (Gopalakrishnan and Winston, 2019) to identify point mutations unique to the suppressor pools, and to monitor ploidy in the suppressor strains. Additional analysis for identification of polymorphisms was performed using Geneious Prime version 2019.0.3 (Kearse et al., 2012). The identified candidate mutations were verified by Sanger sequencing, and the ability of each mutation to confer suppression was verified by genetic tests. These tests (Table S1) included: (a) complementation using plasmids expressing wild-type candidate genes; (b) reconstitution of the suppressor phenotype by null alleles of non-essential genes; and (c) allele replacement of essential genes.

### Mass spectrometry analysis of the RNA polymerase II (RNAPII) interactome

Yeast strains bearing a C-terminal triple FLAG tag on Rpb3 were grown as 1 L cultures to OD_600_ ~ 0.8 in duplicate (FY2912, OV509, OV513) or triplicate (FY3019). Cells were collected by filtration and flash-frozen in liquid nitrogen. After adding 2 ml of frozen buffer A (20 mM Hepes pH 7.6, 20% glycerol, 1 mM DTT, 1 mM EDTA, 125 mM potassium acetate, 1% NP-40), the cells were lysed in a mixer-mill using 8 cycles at 15 Hz. The grindates were thawed on ice in the presence of 4 ml lysis buffer with 1x protease inhibitors (Sigma), and 1x phosphatase inhibitors (Sigma). The lysates were diluted to have equal total protein concentration, and Rpb3 was immunoprecipitated using 200 μl of FLAG M2 beads (Sigma). Samples were then incubated for 2 hours at 4°C on a roller, and the beads were washed three times with buffer A before eluting with FLAG peptide (0.25 mg/ml) in elution buffer (10 mM Tris pH 7.4, 150 mM NaCl, 10% glycerol). The eluates were submitted to the Thermo Fisher Center for Multiplexed Proteomics (Harvard Medical School) for Tandem Mass Tag (TMT)-based mass spectrometry analysis (Zhang and Elias, 2017). The samples were precipitated with trichloroacetic acid, digested with trypsin, labelled using TMT11 reagents, and subjected to LS-MS3 analysis on an Orbitrap Fusion mass spectrometer according to the standard workflow. The resulting peptide spectra were searched using the SEQUEST algorithm against a Uniprot composite database derived from the *S. cerevisiae* proteome containing its reverse complement and known contaminants. Peptide spectral matches were filtered to a 1% false discovery rate (FDR) using the target-decoy strategy combined with linear discriminant analysis. Proteins were quantified only from peptides with a summed SN threshold of >=100 and MS2 isolation specificity of 0.5. Differential protein abundance analysis was performed for proteins with two or more identified peptides, using the Perseus software platform (Hubner et al., 2010; Hubner and Mann, 2011) as described before (Harlen and Churchman, 2017).

### Chromatin immunoprecipitation sequencing (ChIP-seq)

For Spt6 and Spt16 ChIP-seq studies, yeast strains containing Spt6 fused with the triple FLAG epitope tag (FY3276, FY3277, FY3281, FY3282) or Myc-tagged Spt16 fusions (FY3299-3302) were grown in YPD at 30°C or shifted to 37°C as described above. For Spn1 ChIP-seq, strains containing an N-terminal V5 epitope tag on Spn1 (FY3289, FY3292-3294) were grown in YPD at 30°C. The cultures were processed for cross-linking and collecting as previously described (Doris et al., 2018). Chromatin was prepared using standard methods (Gopalakrishnan et al., 2019). Each chromatin sample was mixed with *S. pombe* chromatin (strain FWP570) at 10% level by protein mass for spike-in normalization and split into aliquots of 500 μg of chromatin for immunoprecipitation. Chromatin precipitations were performed overnight at 4°C using either 50 μl of anti-FLAG M2 affinity gel (Sigma) for Spt6, 30 μl of anti-Myc antibodies (9E10, Santa Cruz Biotechnology), 8 μl of 8WG16 antibodies (Millipore Sigma) for Rpb1, or 5 μl of anti-V5 antibodies (Invitrogen) per 500 μg of chromatin. DNA isolated from each sample before precipitation (inputs) and after precipitation (IP) was used for library generation as previously described (Gopalakrishnan et al., 2019).

### Micrococcal nuclease sequencing (MNase-seq)

Cultures (500 ml) for strains FY87, FY3125, FY3223, FY3205, FY3206, FY3272-3274 were grown in YPD at 30°C and also after a shift to 37°C for 80 minutes. The cells were crosslinked with 2% formaldehyde, collected, and subjected to spheroplasting, followed by MNase digestion as previously described (Doris et al., 2018). MNase-digested *S. pombe* DNA (strain 972) was then spiked in according to the original cell count (100 ng of spike-in DNA per 7 × 10^7^ *S. cerevisiae* cells). MNase-seq libraries were generated from the gel-extracted mononucleosomal DNA from each sample as described before (Doris et al., 2018) and were paired-end sequenced on an Illumina NextSeq 500 by the Harvard Bauer Core facility.

## COMPUTATIONAL METHODS

### Data management

All data analyses except for mass spectrometry analyses and polymorphism identification using Geneious were managed using the Snakemake workflow management system (Koster and Rahmann, 2012).

### Genome builds and annotations

The genome builds used were *S. cerevisiae* R64-2-1 (Engel et al., 2014) and *S. pombe* ASM294v2 (Wood et al., 2002). *S. cerevisiae* transcript coordinates were generated from TIF-seq (Pelechano et al., 2013) and TSS-seq data, as previously described (Doris et al., 2018).

### TSS-seq library processing

Removal of adapter sequences and random hexamer sequences from the 3′ end of the read and 3′ quality trimming were performed using cutadapt (Martin, 2011). The random hexamer molecular barcode on the 5′ end of the read was then removed and processed using a Python script. Reads were aligned to the combined *S. cerevisiae* and *S. pombe* reference genomes using Tophat2 (Kim et al., 2013) without a reference transcriptome, and uniquely mapping alignments were selected using SAMtools (Li et al., 2009). Alignments mapping to the same location as another alignment with the same molecular barcode were identified as PCR duplicates and removed using a Python script. Coverage of the 5′-most base, corresponding to the TSS, was extracted using bedtools genomecov (Quinlan and Hall, 2010). Due to high variability in the proportion of *S. pombe* spike-in alignments among the libraries for certain conditions, coverage was normalized to the total number of alignments uniquely mapping to the *S. cerevisiae* genome. The quality of raw, cleaned, non-aligning, and uniquely aligning non-duplicate reads was assessed using FastQC (Andrews, 2014).

### TSS-seq peak calling

TSS peak calling was performed using 1D watershed segmentation followed by filtering for reproducibility by the Irreproducible Discovery Rate method (IDR=0.1) (Li et al., 2011), as previously described in (Doris et al., 2018), except using the maximum signal within a putative peak to estimate the probability of the peak being generated by noise rather than the sum of the signal within the peak. To allow for direct comparison between *spt6-YW* and *spt6-1004,* unified TSS peak sets were generated by using bedtools multiinter to combine peaks called in wild-type, *spt6-YW,* and *spt6-1004* at 30°C or 37°C.

### TSS-seq differential expression analysis

For TSS-seq differential expression analyses, counts of alignments overlapping the unified TSS peak sets described above were used as the input to differential expression analysis by DESeq2 (Love et al., 2014), with a null hypothesis of no change between conditions and a false discovery rate of 0.1.

### Classification of TSS peaks into genomic categories

TSS peak classification was performed as described (Doris et al., 2018), except the summit of intragenic and antisense peaks were used to determine overlap with open/closed reading frames or transcripts. In brief, a genic region was defined for each gene using its transcript and open/closed reading frame annotations. TSS peaks were classified as genic if they overlapped a genic region on the same strand. TSS peaks were classified as intragenic if they were not classified as genic and their summit overlapped an open or closed reading frame on the same strand. TSS peaks were classified as antisense if they were not classified as genic and their summit overlapped a transcript on the opposite strand. TSS peaks not overlapping genic regions, transcripts, or reading frames were classified as intragenic.

### NET-seq library processing

Removal of adapter sequences from the 3′ end of the read and 3′ quality trimming were performed using cutadapt (Martin, 2011). Reads were aligned to the *S. cerevisiae* genome using Tophat2 without a reference transcriptome (kim2013), and uniquely mapping alignments were selected using SAMtools (Li et al., 2009). Coverage of the 5′-most base of the read, corresponding to the 3′-most base of the nascent RNA and the active site of elongating RNA polymerase, was extracted using bedtools genomecov (Quinlan and Hall, 2010), and normalized to the total number of uniquely mapping alignments. The quality of raw, cleaned, non-aligning, and uniquely aligning reads was assessed using FastQC (Andrews, 2014).

### ChIP-seq library processing

Reads were demultiplexed using fastq-multx (Aronesty, 2103), allowing one mismatch to the index sequence and A-tail. Cutadapt (Martin, 2011) was then used to remove index sequences and low-quality base calls from the 3′ end of the read. Reads were aligned to the combined *S. cerevisiae* and *S. pombe* genome using Bowtie 2 (Langmead and Salzberg, 2012), and alignments with a mapping quality of at least 5 were selected using SAMtools (Li et al., 2009). The median fragment size estimated by MACS2 (Zhang et al., 2008b) cross-correlation over all samples of a factor was used to generate coverage of fragments and fragment midpoints by extending alignments to the median fragment size or by shifting the 5′ end of alignments by half the median fragment size, respectively. The quality of raw, cleaned, non-aligning, and uniquely aligning reads was assessed using FastQC (Andrews, 2014).

### ChIP-seq normalization

For ChIP-seq coverage from IP samples, spike-in normalization was accomplished by scaling coverage proportionally to the normalization factor N_input, spike-in_ / N_IP, spike-_ in * N_input, experimental_, where N_IP, spike-in_ is the number of *S. pombe* alignments in the IP sample, N_input,_ spike-in is the number of *S. pombe* alignments in the corresponding input sample, and N_input,_ experimental is the number of *S. cerevisiae* alignments in the input sample. Coverage from input samples was normalized to N_input, experimental_. Relative estimates of the total abundance of each ChIP target on chromatin were also obtained by multiplying the normalization factor with the number of *S. cerevisiae* alignments in an IP sample.

Coverage of the relative ratio of IP over input was obtained by first smoothing normalized IP and input fragment midpoint coverage using a Gaussian with 20 bp bandwidth, and then taking the ratio. Coverage of the relative ratio of one factor to another (e.g. Spn1 over Rpb1) was obtained as follows: For each factor, coverage of IP over input in each sample was standardized using the genome-wide mean and standard deviation over all samples, weighted such that each condition had equal contribution. Standardized coverage of the normalizing factor was then subtracted from the matched coverage of the factor to be normalized.

### ChIP-seq differential occupancy analysis

For differential occupancy analyses of single factors over verified coding genes, IP and input fragment midpoints overlapping the transcript annotation of these genes were counted using bedtools (Quinlan and Hall, 2010). These counts were used to perform a differential occupancy analysis using DEseq2 (Love et al., 2014), at a false discovery rate of 0.1. The design formula used was a generalized linear model with variables for sample type (IP or input), condition (strain and temperature), and the interaction of sample type with condition. Fold changes were extracted from the coefficients of the interaction of sample type with condition, and represent the change in IP signal between conditions, corrected for change in input signal. To normalize to the spike-in control, size factors obtained from *S. pombe* counts over peaks called with MACS2 (Zhang et al., 2008b) and IDR (Li et al., 2011) were used for each sample.

### MNase-seq library processing

Paired-end reads were demultiplexed using fastq-multx (Aronesty, 2103) allowing one mismatch to the barcode. Filtering for the barcode on read 2 and 3′ quality trimming were performed with cutadapt (Martin, 2011). Reads were aligned to the combined *S. cerevisiae* and *S. pombe* genome using Bowtie 1 (Langmead et al., 2009), and correctly paired alignments were selected using SAMtools (Li et al., 2009). Coverage of nucleosome protection and nucleosome dyads were extracted using bedtools (Quinlan and Hall, 2010) and shell scripts to get the entire fragment or the midpoint of the fragment, respectively. Smoothed nucleosome dyad coverage was generated by smoothing dyad coverage was generated by smoothing dyad coverage with a Gaussian kernel of 20 bp bandwidth. Due to differences in the proportion of *S. pombe* DNA added between sequencing runs, coverage was normalized to the total number of correctly paired *S. cerevisiae* fragments. The quality of raw, cleaned, non-aligning, and correctly paired reads was assessed using FastQC (Andrews, 2014).

### MNase-seq quantification

Nucleosome regions for each condition were called using DANPOS2 (Chen et al., 2013). Nucleosome ‘fuzziness’ was calculated for each nucleosome region in each sample by taking the standard deviation of nucleosome dyad positions in the region.

## Notes

### Competing Interest Statement

The authors have declared no competing interest.

