## Supplemental Figures and Tables for Viktorovskaya_2020 for "Essential histone chaperones collaborate to regulate transcription and chromatin integrity"

### Figure S1 (related to Figure 1)

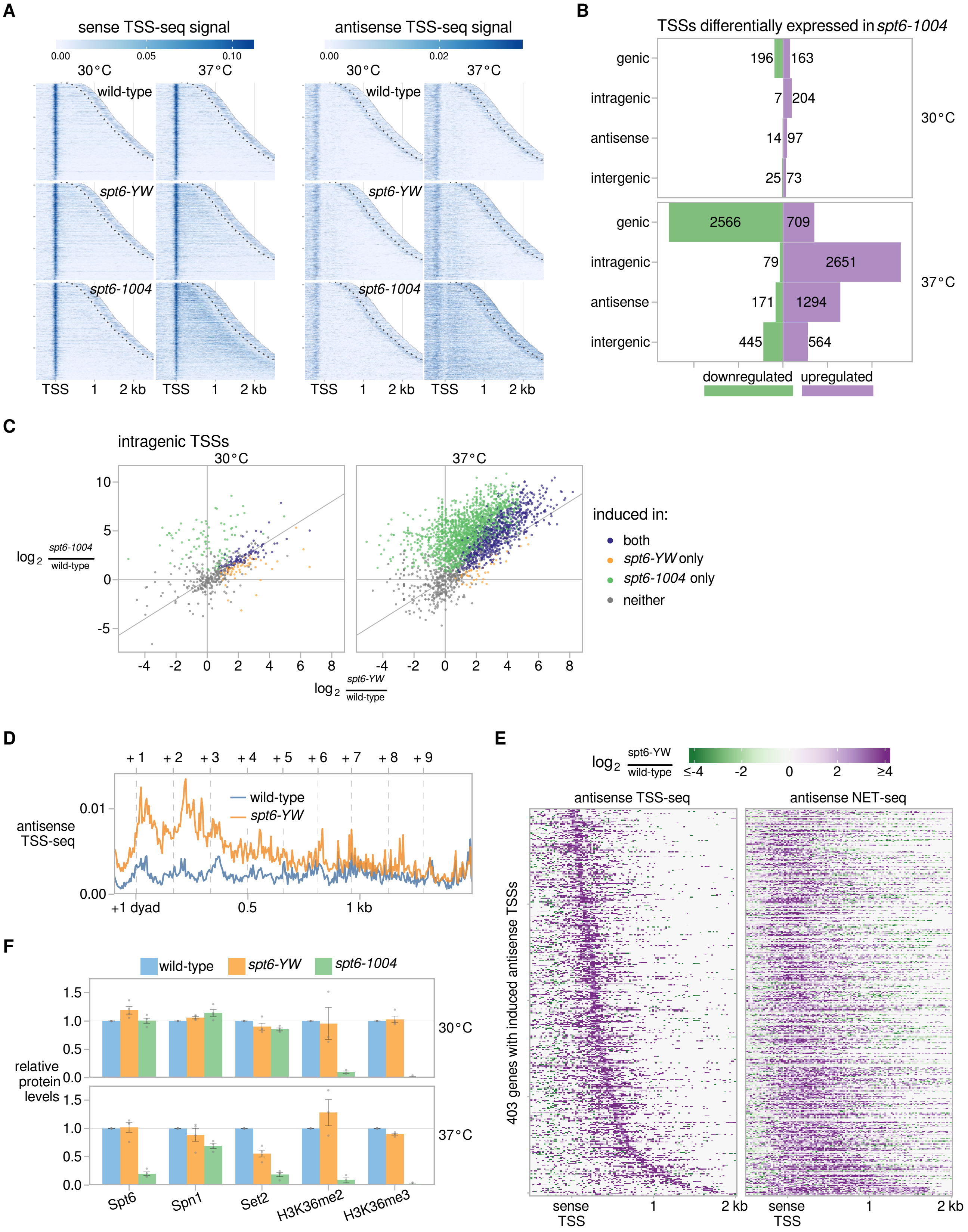

**Figure S1 (related to Figure 1). The *spt6-YW* mutation causes altered sense and antisense transcription**

**(A)** Heatmaps of sense and antisense TSS-seq signal in wild-type, *spt6-YW*, and *spt6-1004* strains, grown either at 30 °C or shifted to 37 °C for 80 minutes. Data are shown for 3,087 non-overlapping verified coding genes aligned by wild-type genic TSS and sorted by length, up to 300 nt 3′ of the cleavage and polyadenylation site indicated by the white dotted line.

**(B)** Bar plots showing the number of TSS-seq peaks differentially expressed in *spt6-1004* versus wild-type, separated by genomic class.

**(C)** Scatterplots showing changes in sense strand intragenic transcript abundance versus wild-type strains, comparing *spt6-1004* to *spt6-YW*. TSS peaks are colored based on significant upregulation in one, both, or neither mutant.

**(D)** Median antisense TSS-seq signal in wild-type and *spt6-YW* at 37 °C, over 3,086 non-overlapping verified coding genes aligned by wild-type +1 nucleosome dyad. The average positions of the wild-type +1 through +9 nucleosome dyads as determined from MNase-seq data are indicated with vertical dashed lines.

**(E)** Heatmaps of change in antisense TSS-seq and NET-seq signal in *spt6-YW* versus wild-type at 37 °C. Data are shown over 403 genes with a significantly induced antisense TSS peak in *spt6-YW* at 37 °C, aligned by wild-type sense TSS and arranged by the distance from the sense TSS to the *spt6-YW*-induced antisense TSS.

**(F)** Quantification of western blots for protein abundance in whole cell lysates of wild-type, *spt6-YW*, and *spt6-1004* strains grown at 30 °C or shifted to 37 °C for 80 minutes. Spt6, Spn1, and Set2 signal were normalized to a Pgk1 loading control, and histone modification signal was normalized to histone H3. Error bars indicate the mean ± standard error of the replicates shown. Samples from a *set2Δ* strain were included as a negative control for the histone modifications (data not shown).

### Figure S2 (related to Figure 3)

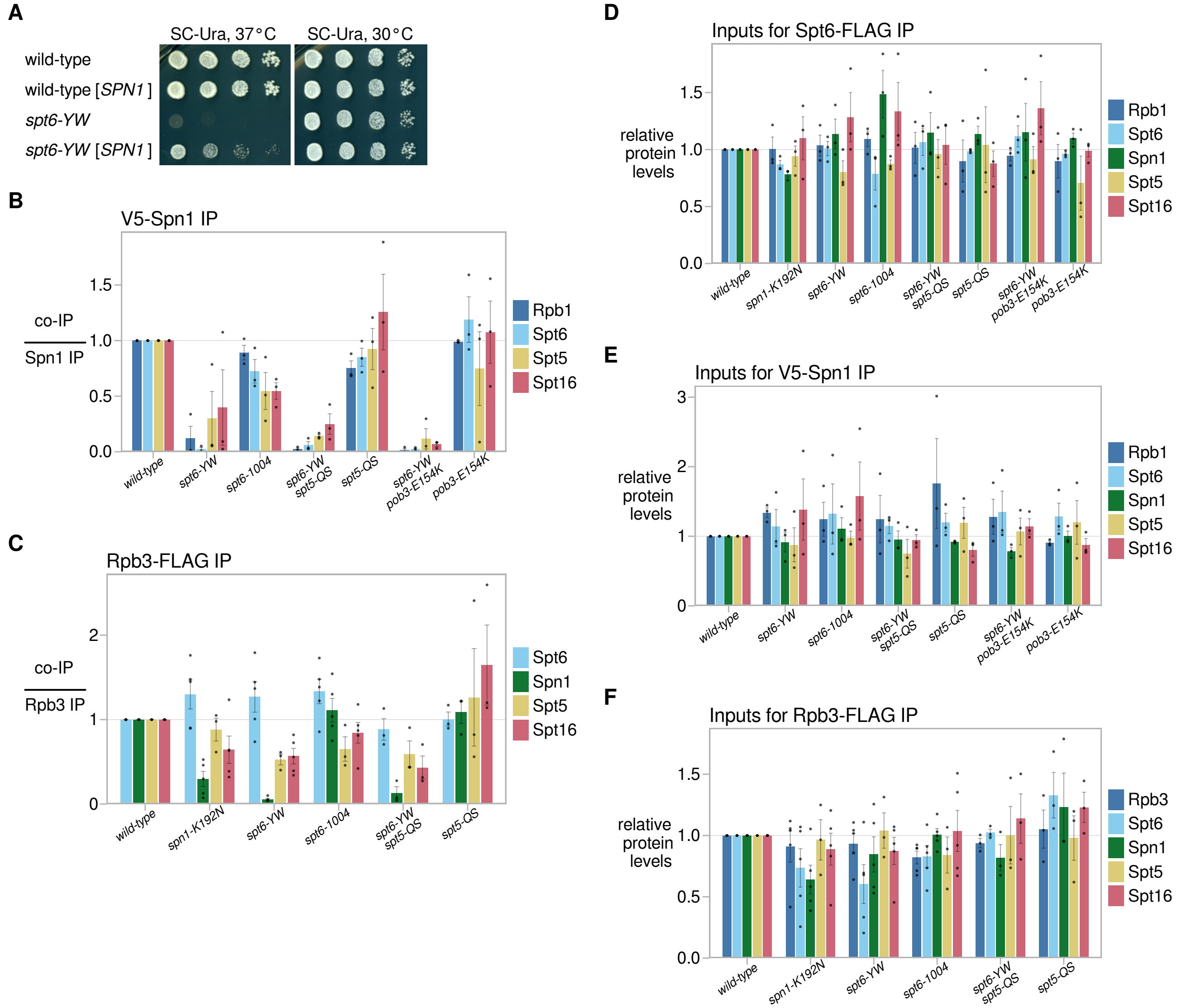


**Figure S2 (related to Figure 3). The *pob3-E154K* and *spt5-QS* suppressors do not restore Spt6-Spn1 interaction**

**(A)** Analysis of effects of *SPN1* overexpression on *spt6-YW* temperature sensitivity. Strains FY3276 and FY3277, transformed with either plasmid FB2701 [*SPN1-Myc*, *CEN*, *URA3*, *HIS3*] or an empty vector (see Key Resources Table), were grown to saturation in liquid media without uracil, serially diluted 10-fold, spotted on the indicated media, and grown at the indicated temperature.

**(B)** Quantification of Spn1-V5 co-immunoprecipitation experiments. Error bars indicate the mean ± standard error of the relative co-immunoprecipitation signal normalized to the Spn1-V5 pull-down signal in the replicates shown.

**(C)** Quantification of Rpb3-FLAG co-immunoprecipitation experiments. Error bars indicate the mean ± standard error of the relative co-immunoprecipitation signal normalized to the Rpb3-FLAG pull-down signal in the replicates shown.

**(D)** Quantification of protein abundance from western blots of the inputs for Spt6-FLAG co-immunoprecipitation experiments, as shown in Figure 3A. Error bars indicate the mean ± standard error of the relative western blot signal from the replicates shown.

**(E)** As in (D), but for V5-Spn1 inputs.

**(F)** As in (D), but for Rpb3-FLAG inputs.

### Figure S3 (related to Figure 4)

**A**

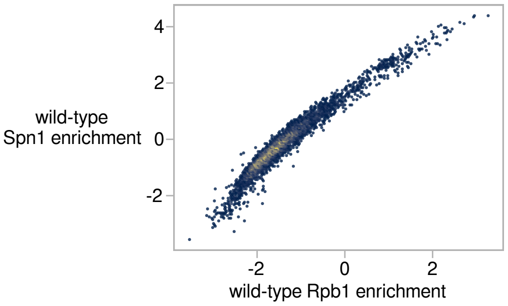


**B**

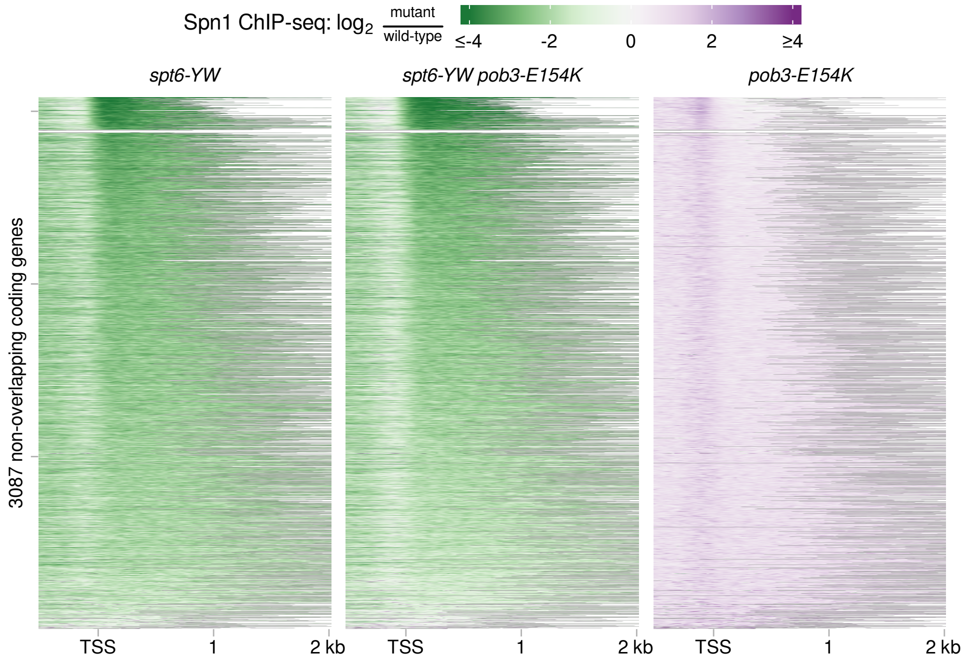


**Figure S3 (related to Figure 4). The Spt6-dependent recruitment of Spn1 to chromatin is not altered by *pob3-E154K***

**(A)** Scatterplot of Spn1 ChIP enrichment versus Rpb1 enrichment for 5,091 verified coding genes in wild-type cells. Enrichment values are the relative $log_2$ enrichment of IP over input.

**(B)** Heatmaps of the change in Spn1 enrichment in mutants over wild-type for 3,087 non-overlapping verified coding genes aligned by TSS and arranged top to bottom by decreasing wild-type Rpb1 enrichment.

### Figure S4 (related to Figure 5)

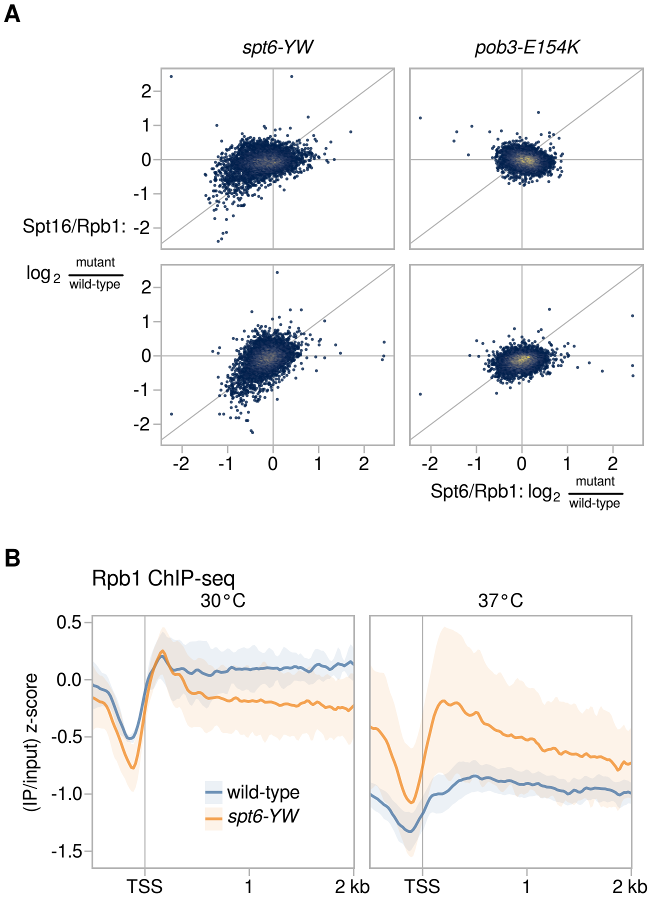


**Figure S4 (related to Figure 5). Evidence that mutant FACT suppresses the Spt6-Spn1 defect by restoring the balance between FACT and Spt6 on chromatin**

**(A)** Scatterplots showing changes in Rpb1-normalized Spt16 ChIP enrichment in mutants over wild-type versus changes in Rpb1-normalized Spt6 ChIP enrichment, for 5,091 verified coding genes.

**(B)** Average Rpb1 ChIP enrichment over 3,087 non-overlapping verified coding genes aligned by TSS in wild-type (FY3276) and *spt6-YW* (FY3277) strains, grown either at 30°C or with an 80-minute shift to 37°C. For each gene, the spike-in normalized ratio of IP over input signal is standardized to the mean and standard deviation of the 30°C wild-type signal over the region. The solid line and shading are the mean and 95% confidence interval of the mean standard score over the genes considered from two to five replicates.

### Figure S5 (related to Figure 6)

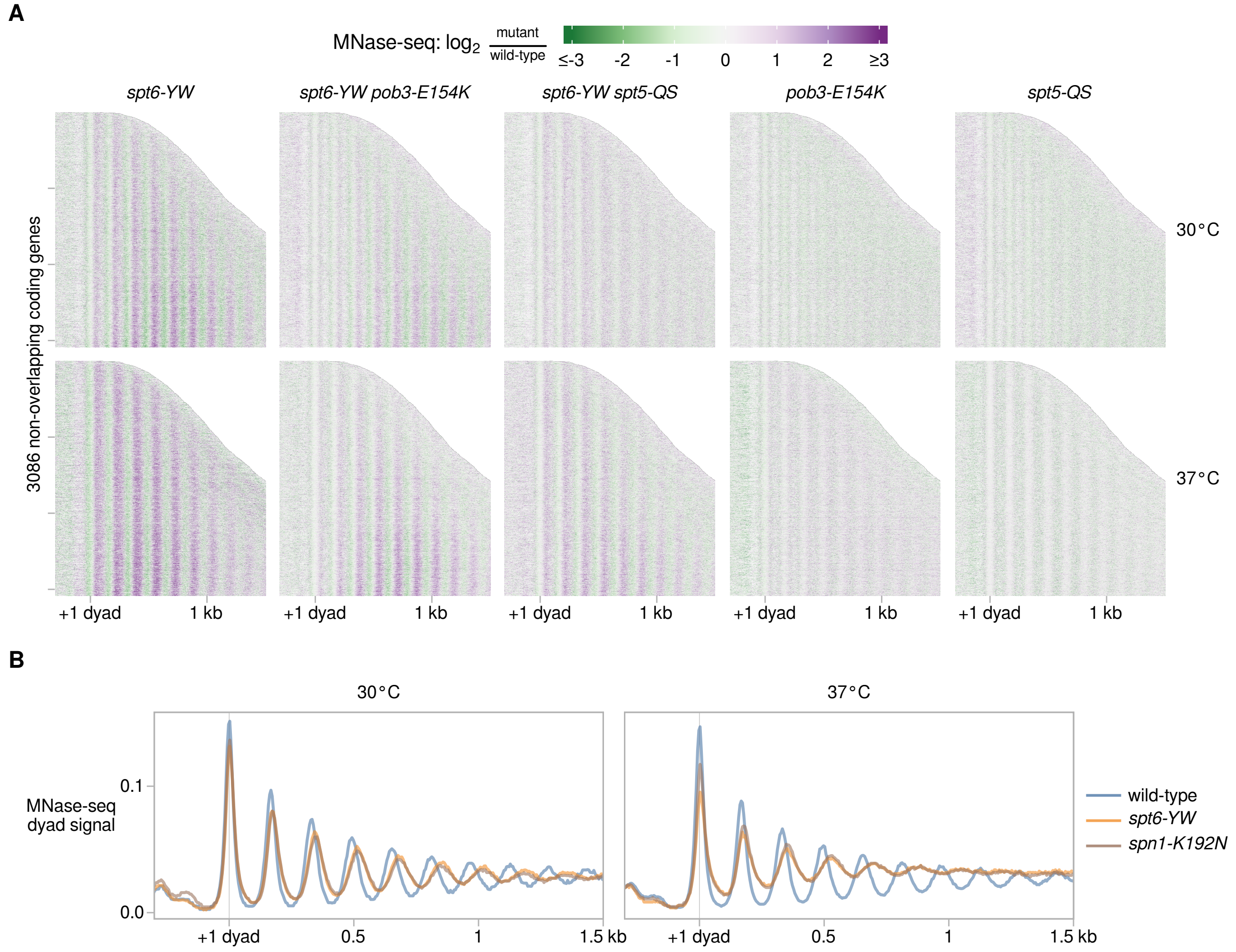


**Figure S5 (related to Figure 6). Spt6, FACT, Spt5, and Spn1 functionally interact to modulate nucleosome organization *in vivo***

**(A)** Heatmaps of changes to MNase-seq dyad signal in mutants over wild-type for 3086 non-overlapping coding genes aligned by wild-type +1 nucleosome dyad and sorted by gene length, for strains grown either at 30°C or with an 80-minute shift to 37°C.

**(B)** Average MNase-seq dyad signal over 3,086 non-overlapping verified coding genes aligned by 30°C wild-type +1 nucleosome dyad, for wild-type (FY87), *spt6-YW* (FY3223), *spn1-K192N* (FY3272) strains grown at 30°C or with an 80-minute shift to 37°C. Values are the mean of the mean library-size normalized coverage over the genes considered, over at least two replicates.

**Table S1 (related to Table 1). Identification and genetic verification of the suppressors for *spt6-YW* temperature-sensitive phenotype.**

| Gene | Mutation or amino acid change | Times isolated | Genetic verification | | |
|---|---|---|---|---|---|
| | | | Phenotype linkage[1] | Complementation by respective WT[2] | Allele reconstitution[3] |
| Intragenic to *SPT6* | | | | | |
| *SPT6* | **P231L**, Y255A, W257A | 3 | Linked to *SPT6* | ND | ND |
| Extragenic | | | | | |
| *HTA1* | I80S | 1 | Linked to *HTA1* | phenotype rescue by *HTA1* | ND |
| *RCO1* | S558X | 1 | Linked to *RCO1* | phenotype rescue by *RCO1* | *rco1Δ* suppresses *spt6-YW* |
| | C440X | 1 | | | |
| *SET2* | K213E G219K | 1 | Linked to *SET2* | phenotype rescue by *SET2* | *set2Δ* suppresses *spt6-YW* |
| *CHD1* | frameshift after codon 812 | 1 | Linked to *CHD1* | ND | *chd1Δ* suppresses *spt6-YW* |
| *SPT5* | Q342H S343del (3 bp deletion) | 1 | Linked to *SPT5* | phenotype rescue by *SPT5* | *spt5-QS* suppresses *spt6-YW* |
| *POB3* | E154K | 7 | Linked to *POB3* | phenotype rescue by *POB3* | *pob3-E154K* suppresses *spt6-YW* |
| | P253L | 1 | | | |
| *SPT16* | T627K | 1 | Linked to *SPT16* | phenotype rescue by *SPT16* | ND |
| | E656K | 1 | | | |
| | K579E | 1 | | | |
| **Disomic** | | | | | |
| chr XVI disomy[4] | | 8 | NA | NA | *SPN1* on CEN plasmid suppresses *spt6-YW* |

[1] Phenotype linkage was assessed by the analysis of the $Ts^+$ phenotypes in the progeny segregating the indicated suppressor allele in the *spt6-YW* backgrounds.

[2] Complementation was examined by transforming the indicated *spt6-YW* suppressor strains with a plasmid expressing a respective wild-type allele for the suppressor gene (see Key Resource Table) followed by the phenotype analysis of the transformants. Phenotype rescue was scored when transformants had a $Ts^-$ phenotype (complete rescue), similar to the *spt6-YW* parent strains, or had an intermediate $Ts^{\pm}$ phenotype (partial rescue). "ND" stands for "not determined".

[3] In case of the non-essential genes (*RCO1*, *SET2*, and *CHD1*), the null alleles were shown to suppress the Ts- phenotype of *spt6-YW* (Table S2). In case of FACT and Spt5, the identified mutations were introduced to *SPT5* and *POB3* genes, reconstituting the *spt5-QS* and *pob3-E154K* alleles, and genetic suppression of *spt6-YW* by either allele was confirmed (Figure 2B). "ND" stands for "not determined".

[4] In several cases the chromosome XVI disomy was accompanied by other disomies, including chr.I, III or XI. None of the disomies other than the chr.XVI disomy were specifically associated with the $Ts^+$ suppressor phenotype.

**Table S2 (related to Figure 2). The phenotypes for different mutant combinations used to determine genetic interactions.**

| Strain | 30°C YPD | 37°C YPD | 25°C YPD | Spt[1] (SC -Lys) | Phleomycin[2] | Hydroxyurea[3] |
|---|---|---|---|---|---|---|
| wild type | ++++ | ++++ | ++++ | - | +++ | ++++ |
| *pob3-E154K* | ++++ | ++++ | ++++ | - | +++ | ++++ |
| *spt5-QS* | ++++ | ++++ | ++++ | - | ++ | ++++ |
| *set2Δ* | ++++ | ++++ | ++++ | - | +++ | ++++ |
| *rco1Δ* | ++++ | ++++ | ++++ | - | ND | ND |
| *chd1Δ* | ++++ | ++++ | ++++ | ND | ND | ++++ |
| *pob3-272 (I282K)* | +++ | +++ | +++ | ++++ | ND | ND |
| *spt16-197 (G132D)* | ++++ | + | ++++ | ++ | ND | ND |
| *spt5-4 (E338K)* | +++ | ++ | ++ | ++ | ND | ND |
| *spt5-194 (S324F)* | +++ | +++ | ND | +++ | ND | ND |
| *spt5-242 (A268V)* | ++++ | ++++ | - | + | ND | ND |
| *spt4Δ* | ++++ | ++ | ND | ND | ND | ND |
| *spt6-YW* | ++++ | - | ++++ | ++++ | + | + |
| *spt6-YW pob3-E154K* | ++++ | ++++ | ++++ | ++ | ++ | +++ |
| *spt6-YW spt5-QS* | ++++ | ++++ | ++++ | ++++ | ++ | +++ |
| *spt6-YW set2Δ* | ++++ | ++ | ++++ | +++ | + | ++ |
| *spt6-YW rco1Δ* | ++++ | ++ | ++++ | ++++ | ND | ND |
| *spt6-YW chd1Δ* | ++++ | ++ | ++++ | ND | ND | +++ |
| *spt6-YW pob3-272* | +++ | - | +++ | +++ | ND | ND |
| *spt6-YW spt16-197* | ++++ | - | ++++ | +++ | ND | ND |
| *spt6-YW spt5-4* | inviable | | | | | |
| *spt6-YW spt5-194* | inviable | | | | | |
| *spt6-YW spt5-242* | ++++ | - | +++ | ++++ | ND | ND |
| *spt6-YW spt4Δ* | inviable | | | | | |
| *spt6-F249K* | ++++ | - | ++++ | ++ | ND | ND |
| *spt6-F249K pob3-E154K* | ++++ | ++++ | ++++ | - | ND | ND |
| *spt6-F249K spt5-QS* | ++++ | ++++ | ++++ | ++++ | ND | ND |
| *spt6-F249K rco1Δ* | ++++ | +++ | ++++ | ++++ | ND | ND |
| *spt6-1004* | ++++ | + | ++++ | ++++ | ND | ND |
| *spt6-1004 pob3-E154K* | ++++ | - | ++++ | +++ | ND | ND |
| *spt6-1004 spt5-QS* | ++++ | - | ++++ | ++++ | ND | ND |
| *spt6-1004 rco1Δ* | +++ | - | +++ | +++ | ND | ND |
| *spn1-K192N* | ++++ | - | ++++ | ++++ | + | + |
| *spn1-K192N pob3-E154K* | ++++ | ++++ | ++++ | ++++ | +++ | ++++ |
| *spn1-K192N spt5-QS* | ++++ | ++++ | ++++ | ++++ | + | ++++ |
| *spn1-K192N chd1Δ* | ++++ | +[4] | ++++ | ND | ND | +++ |
| *spt6-YW spn1-K192N* | inviable | | | | | |
| *spt6-YW spn1-K192N pob3-E154K* | ++++ | ++ | ++++ | +++ | ND | ND |
| *spt6-YW spn1-K192N spt5-QS* | inviable | | | | | |
| *spn1Δ* | inviable | | | | | |
| *spn1Δ pob3-E154K* | +++ | ++ | ND | +++ | ND | ND |
| *spn1Δ spt5-QS* | inviable | | | | | |

The phenotypes were analyzed based on the yeast growth under indicated conditions in comparison to the wild-type strain using the spot test assay and scored after two days of incubation, unless indicated otherwise. The number of "+" indicates growth of each of the 10-fold serially diluted cultures. "-" indicates a very weak growth, if any. "ND" stands for "not determined".

[1] Spt phenotype was scored as growth on synthetic media without lysine due to suppression of the *lys2-128δ* allele.

[2] Phleomycin was added to the YPD media at concentration of 13 μg/ml; plates were incubated at 30°C for three days before scoring.

[3] Hydroxyurea was supplemented to YPD at 150 mM final concentration; plates were incubated at 30°C.

[4] *spn1-K192N chd1Δ* mutant was scored for growth at 37°C after 3 days of incubation, indicating weak suppression of Ts- phenotype.

**Table S4. The list of strains used in the study.**

| Strain | Genotype | Used for |
|---|---|---|
| *S. cerevisiae* | | |
| FY87 | *MATα lys2-128δ ura3-52 leu2Δ1* | Genetics, TSS-seq, MNase-seq, |
| FY3223 | *MATα spt6-YW lys2-128δ ura3-52 leu2Δ1 [pRS316]* | |
| FY3207 | *MATα spt6-YW lys2-128δ ura3-52 leu2Δ1* | |
| FY3205 | *MATα spt6-YW pob3-E154K lys2-128δ ura3-52 leu2Δ1 [pRS316]* | |
| FY3206 | *MATα pob3-E154K ura3-52 lys2-128δ [pRS316]* | |
| FY3125 | *MATα spt6-1004 lys2-128δ ura3-52 leu2Δ1* | |
| FY3272 | *MATα spn1-K192N::URA3 leu2D0 lys2-128δ ura3D0* | |
| FY3273 | *MATα spt5-QS lys2-128δ ura3-52 leu2Δ1* | |
| FY3274 | *MATα spt5-QS spt6-YW lys2-128δ ura3-52 leu2Δ1* | |
| FY3220 | *MATα spt6-YW pob3-E154K lys2-128δ ura3-52 leu2Δ1* | |
| FY3221 | *MATα pob3-E154K ura3-52 leu2Δ1 lys2-128δ* | |
| FY3276 | *MATα SPT6-(FLAG)*$_{X3}$ *lys2-128δ ura3-52 leu2Δ1* | Spt6 ChIP-seq Spt6 pulldown |
| FY3277 | *MATα spt6-YW-(FLAG)x3 lys2-128δ ura3-52 leu2Δ1* | |
| FY3278 | *MATα spt6-YW-(FLAG)x3 spt5-QS ura3-52 leu2Δ1 lys2-128δ* | |
| FY3279 | *MATα SPT6-(FLAG)x3 spt5-QS ura3-52 leu2Δ1 lys2-128δ* | |
| FY3280 | *MATα SPT6-(FLAG)x3 spn1-K192N::URA3 spt5-QS ura3-52 leu2Δ1 lys2-128δ* | |
| FY3281 | *MATα spt6-YW-(FLAG)x3 pob3-E153K ura3-52 leu2Δ1 lys2-128δ* | |
| FY3282 | *MATα SPT6-(FLAG)x3 pob3-E153K ura3-52 leu2Δ1 lys2-128δ* | |
| FY3283 | *MATα spt6-1004-(FLAG)x3 lys2-128δ ura3-52 leu2Δ1* | |
| FY2912 | *MATα RPB3-(FLAG)x3::NatMx ura3-52 his4-912δ lys2-128δ* | NET-seq Rpb3 pulldown and/or IP-MS |
| FY3019 | *MATα RPB3-(FLAG)x3::NatMx spt6-YW his4-912δ lys2-128δ ura3-52* | |
| FY3021 | *MATα RPB3-(FLAG)x3::NatMx spt6-1004 his4-912δ lys2-128δ ura3-52* | |
| FY3284 | *MATα RPB3-(FLAG)x3::NatMx spt6-YW spt5-QS his4-912δ lys2-128δ ura3-52* | |
| FY3285 | *MATα RPB3-(FLAG)x3::NatMx spt5-QS his4-912δ lys2-128δ ura3-52* | |
| FY3286 | *MATα RPB3-(FLAG)x3::NatMx spn1-K192N::URA3 his4-912δ lys2-128δ ura3-52* | |
| FY3287 | *MATα RPB3-(FLAG)x3::NatMx spt6-YW pob3-E154K his4-912δ lys2-128δ ura3-52* | |
| FY3288 | *MATα RPB3-(FLAG)x3::NatMx pob3-E154K his4-912δ lys2-128δ ura3-52* | |
| FY3289 | *MATα (V5)x3-SPN1 spt6-YW ura3-52 lys2-128δ leu2Δ1 his3Δ200* | Spn1 ChIP-seq Spn1 pulldown |
| FY3290 | *MATα (V5)x3-SPN1 spt6-YW spt5-QS ura3-52 lys2-128δ leu2Δ1 his3Δ200* | |
| FY3291 | *MATα (V5)x3-SPN1 spt5-QS ura3-52 lys2-128δ leu2Δ1 his3Δ200* | |
| FY3292 | *MATα (V5)x3-SPN1 ura3-52 lys2-128δ leu2Δ1 his3Δ200* | |
| FY3293 | *MATα (V5)x3-SPN1 pob3-E154K spt6-YW ura3-52 lys2-128δ leu2Δ1 his3Δ200* | |
| FY3294 | *MATα (V5)x3-SPN1 pob3-E154K ura3-52 lys2-128δ leu2Δ1 his3Δ200* | |
| FY3296 | *MATα (V5)x3-SPN1 spt6-1004 ura3-52 lys2-128δ leu2Δ1 his3Δ200* | |
| FY3297 | *MAT*__a__ *spt6-YW his4-912δ lys2-128δ ura3-52 leu2Δ1* | suppressor isolation |
| FY3298 | *MATα spt6-YW his4-912δ lys2-128δ ura3-52* | |
| FY3299 | *MATα SPT16-Myc leu2Δ1 ura3-52 lys2-128δ* | Spt16 ChIP-seq |
| FY3300 | *MATα SPT16-Myc spt6-YW pob3-E154K leu2Δ1 ura3-52 lys2-128δ* | |
| FY3301 | *MATα SPT16-Myc spt6-YW leu2Δ1 ura3-52 lys2-128δ* | |
| FY3302 | *MATα SPT16-Myc pob3-E154K leu2Δ1 ura3-52 lys2-128δ* | |
| FY3303 | *MAT*__a__ *pob3-E154K-(V5)x3 spt6-YW lys2-128δ ura3-52 leu2Δ1* | Pob3 ChIP Pob3 pulldown |
| FY3304 | *MAT*__a__ *POB3-(V5)x3 lys2-128δ ura3-52 leu2Δ1* | |
| FY3305 | *MAT*__a__ *POB3-(V5)x3 spt6-YW lys2-128δ ura3-52 leu2Δ1* | |
| FY3306 | *MAT*__a__ *pob3-E154K-(V5)x3 lys2-128δ ura3-52 leu2Δ1* | |

| | | |
|---|---|---|
| FY3307 | *MAT***a** *set2Δ::KanMX his3D200 lys2-128δ ura3-52 leu2Δ1 FLO8-URA3* | |
| FY3308 | *MATα rco1Δ::KanMX ura3-52 his4-912δ lys2-128δ* | |
| FY3309 | *MAT***a** *chd1D::hphMX ura3-52 his4-912δ leu2D0 lys2-128δ* | |
| O877 | *MATα pob3-272 his4-912δ lys2-128δ leu2Δ1 ura3-52 suc2dUAS* | |
| FY346 | *MAT***a** *spt16-197 ura3-52 leu2Δ1 lys2-128δ* | |
| FY1668 | *MAT***a** *spt5-4 his4-912δ lys2-128δ* | |
| FY300 | *MAT***a** *spt5-194 his4-912δ lys2-128δ ura3-52 leu2Δ1* | |
| FY1672 | *MATα spt5-242 lys2-128δ leu2Δ1 ura3-52* | |
| FY247 | *MAT***a** *spt4Δ::URA3 ura3-52 leu2Δ1* | |
| FY3310 | *MATα set2Δ::kanMX6 spt6-YW ura3-52 lys2-128δ* | |
| FY3311 | *MAT***a** *spt6-YW rco1Δ::KanMX ura3-52 lys2-128δ his4-912δ* | |
| FY3312 | *MATα spt6-YW chd1Δ::hphMX lys2-128δ ura3-52 his4-912δ leu2Δ1* | |
| FY3313 | *MATα spt6-YW pob3-272 ura3-52 lys2-128δ his4-912δ leu2Δ1 suc2dUAS* | |
| FY3314 | *MAT***a** *spt6-YW spt16-197 lys2-128δ his4-912δ* | |
| FY3315 | *MATα spt6-YW spt5-242 lys2-128δ his4-912δ leu2Δ1* | Genetic Interactions |
| FY3316 | *MATα spt6-F249K(-424, URA3) lys2-128δ ura3-52 his4-921δ* | |
| FY3317 | *MATα spt6-F249K(-424, URA3) pob3-E154K lys2-128δ ura3-52 leu2Δ1 his4-921δ* | |
| FY3318 | *MATα spt6-F249K(-424, URA3) spt5-QS lys2-128δ ura3-52* | |
| FY3319 | *MATα spt6-F249K(-424, URA3) rco1Δ::KanMx his4-912δ lys2-128δ ura3-52* | |
| FY3320 | *MATα spt6-1004 pob3-E154K ura3-52 lys2-128δ leu2Δ1* | |
| FY3321 | *MAT***a** *spt6-1004 spt5-QS lys2-128δ ura3-52 leu2Δ1* | |
| FY3322 | *MATα spt6-1004 rco1Δ::KanMx his4-912δ lys2-128δ ura3-52* | |
| FY3323 | *MATα spn1-K192N::URA3 pob3-E154K lys2-128δ ura3 leu2Δ1* | |
| FY3324 | *MATα spn1-K192N::URA3 spt5-QS lys2-128δ ura3 leu2D0* | |
| FY3325 | *MATα spn1-K192N::URA3 chd1Δ::hphMX ura3-52 leu2D0 lys2-128δ* | |
| FY3326 | *MATα spt6-YW spn1-K192N::URA3 pob3-E154K ura3 lys2-128δ* | |
| FY3327 | *MATα spn1Δ::KanMX his3d200 leu2Δ1 ura3-52 lys2-128δ [SPN1, URA3]* | |
| FY3328 | *MATα spn1Δ::KanMX pob3-E154K his3d200 lys2-128δ ura3-52 [SPN1, URA3]* | |
| FY3329 | *MATα spn1Δ::KanMX spt5-QS his3d200 lys2-128δ ura3-52 [SPN1, URA3]* | |
| *S. pombe* | | |
| FWP570 | *h+ spt5::spt5-V5-IAA::KanMx rpb3-3XFLAG::ura4+ ctr9-Myc::KanMx ura4-D18 ade6-210* | spike-in for ChIP-seq |
| 972 | *h- wild-type* | spike-in for TSS-seq and MNase-seq |
